# Autism-linked Cullin3 germline haploinsufficiency impacts cytoskeletal dynamics and cortical neurogenesis through RhoA signaling

**DOI:** 10.1101/2020.02.07.939256

**Authors:** Megha Amar, Akula Bala Pramod, Nam-Kyung Yu, Victor Munive Herrera, Lily R. Qiu, Patricia Moran-Losada, Pan Zhang, Cleber A. Trujillo, Jacob Ellegood, Jorge Urresti, Kevin Chau, Jolene Diedrich, Jiaye Chen, Jessica Gutierrez, Jonathan Sebat, Dhakshin Ramanathan, Jason P. Lerch, John R. Yates, Alysson R. Muotri, Lilia M. Iakoucheva

**Affiliations:** Department of Psychiatry, University of California San Diego, La Jolla, CA, USA; Department of Molecular Medicine, The Scripps Research Institute, La Jolla, CA, USA; Mouse Imaging Centre (MICe), Hospital for Sick Children, Toronto, ON, Canada; Department of Cellular & Molecular Medicine, University of California San Diego, La Jolla, CA USA; Department of Pediatrics/Rady Children’s Hospital San Diego, University of California San Diego, La Jolla, CA, USA; Beyster Center for Psychiatric Genomics, University of California San Diego, La Jolla, CA, USA; Kavli Institute for Brain and Mind, University of California San Diego, La Jolla, CA, USA; Center for Academic Research and Training in Anthropogeny (CARTA), La Jolla, CA, USA

**Keywords:** Cullin 3 ubiquitin ligase, autism spectrum disorder, Cul3 haploinsufficiency, mouse model, actin cytoskeleton, intermediate filament, microtubule, neurogenesis, dendrite length, early brain development, network activity, Rho signaling, Rhosin rescue

## Abstract

E3-ubiquitin ligase Cullin3 (*Cul3*) is a high confidence risk gene for Autism Spectrum Disorder (ASD) and Developmental Delay (DD). To investigate how *Cul3* mutations impact brain development, we generated haploinsufficient *Cul3* mouse model using CRISPR/Cas9 genome engineering. *Cul3* mutant mice exhibited social and cognitive deficits and hyperactive behavior. Brain MRI found decreased volume of cortical regions and changes in many other brain regions of *Cul3* mutant mice starting from early postnatal development. Spatiotemporal transcriptomic and proteomic profiling of the brain implicated neurogenesis and cytoskeletal defects as key drivers of *Cul3* functional impact. Specifically, dendritic growth, filamentous actin puncta, and spontaneous network activity were reduced in *Cul3* mutant mice. Inhibition of small GTPase RhoA, a molecular substrate of Cul3 ligase, rescued dendrite length and network activity phenotypes. Our study identified neuronal cytoskeleton and Rho signaling as primary targets of *Cul3* mutation during early brain development.

## Introduction

Rare and *de novo* single nucleotide variants (SNVs) and copy number variants (CNVs) are major risk factors for neurodevelopmental disorders (NDDs). E3 ubiquitin ligase Cullin 3 (*Cul3*) is among genes most confidently implicated in NDDs with the genome-wide significant FDR<0.01 ^1^. Exome sequencing studies have identified at least 20 mutations in the *Cul3* gene (13 protein-truncating mutations and 7 missense) in patients with Autism Spectrum Disorder (ASD), Developmental Delay (DD) and Schizophrenia (SCZ), with no *Cul3* mutations detected in healthy controls (**Figure 1a**, **Supplementary Table S1**).

**Figure 1.**
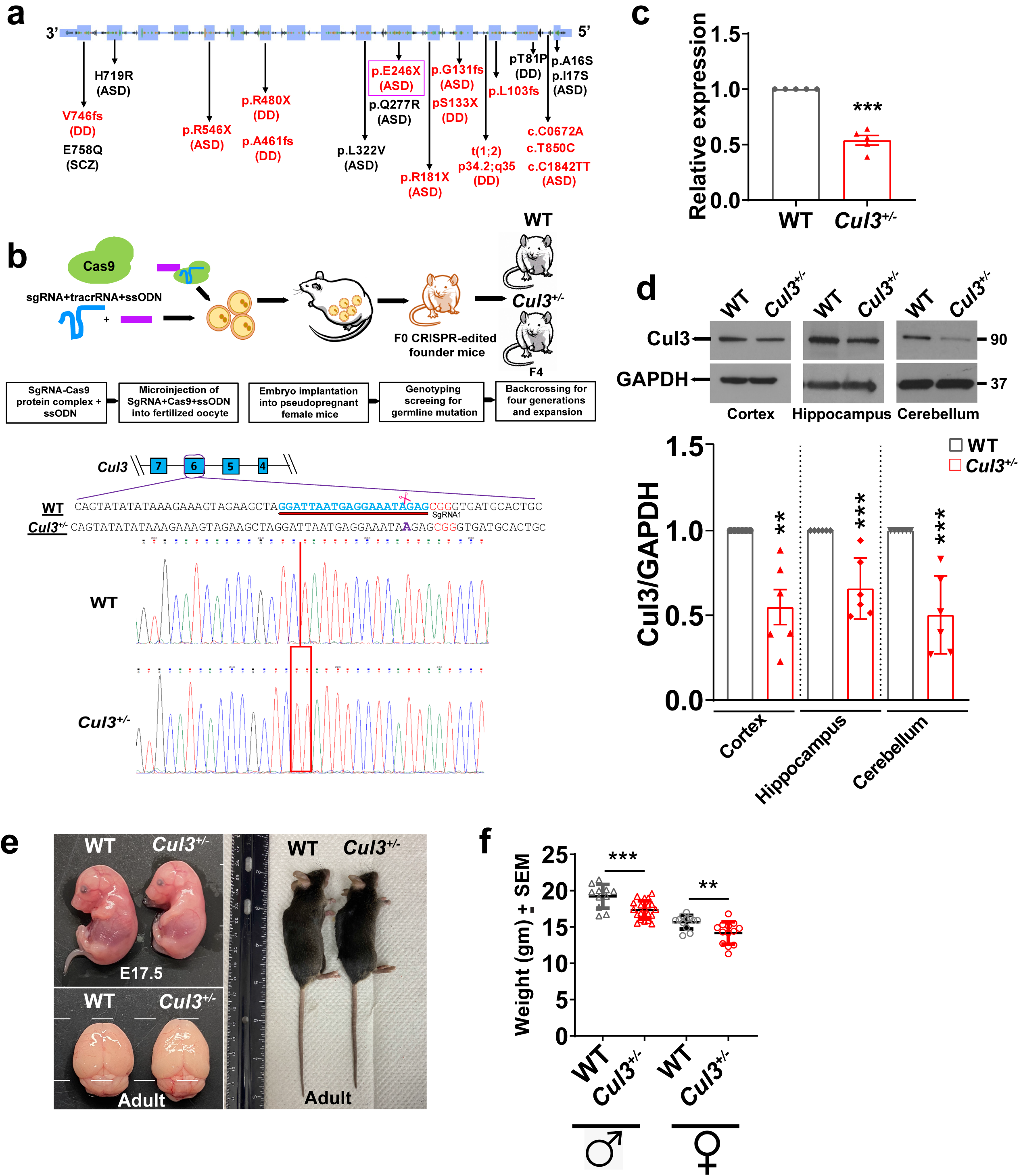
Generation and characterization of *Cul3^+/-^* mouse model. **(a)** Loss-of-function and missense mutations in *Cul3* gene identified in the patients with neurodevelopmental disorders. ASD, autism spectrum disorder; SCZ – schizophrenia; DD – developmental delay. The location of E246X patient’s mutation from this study is highlighted in magenta box. (**b)** Schematic representation of mouse line generation using CRISPR/Cas9 genome editing. cDNA sequence trace showing 1-bp insertion (purple) in exon 6 is shown. Single guide (Sg) RNA is shown in blue, PAM site is shown in red. Sanger sequencing diagram showing 1bp insertion is at the bottom. **(c)** qRT-PCR showing reduction of mRNA expression of Cul3 in adult cerebral cortex of *Cul3^+/-^* mice (***p<0.001; two tailed T-test; n=5 for both genotypes); Error bars represents mean±SEM. **(d)** Western blot of Cul3 protein, showing its significant reduction in *Cul3^+/-^* mutant mice cortex (p<0.01; n=6 for each genotype); hippocampus (p<0.001; n=6 for each genotype) and cerebellum (p<0.001; n=6 for each genotype); error bars represents mean±SD. **(e)** (top left panel) Representative image of embryonic day E17.5 WT and *Cul3^+/-^* embryo demonstrating smaller size; (bottom left panel) WT and *Cul3^+/-^* adult brain; (right panel) Representative image of adult (8-weeks old) WT and *Cul3^+/-^* males demonstrating smaller body size of *Cul3^+/-^* mutants. **(f)** Both adult male and female *Cul3^+/-^* mice have reduced body weight as compared to their WT littermates (***p<0.001; **p<0.01; Two-tailed T-test) (WT n=11 male/12 female; *Cul3^+/-^* n=21 male/14 female) ○-female and Δ-male; Error bars represents mean±SEM. Two tailed T-test was used for calculating statistical significance for **c-f**. Dots represents individual animals.

Cul3 belongs to a family of RING E3 ubiquitin ligases, and its structure and function has been well-characterized, mostly in non-neuronal cell types ^2^. Cul3 regulates turnover of a variety of substrate proteins by ubiquitinating and directing them for proteasomal degradation. Cul3 can interact with different adaptor proteins, and is involved in regulation of a wide range of cellular processes including cytoskeleton organization, cell fate determination, cell cycle regulation, and stress response among many others ^2^. Less is known about the function of Cul3 in neural cells, despite its increased expression during embryonic stages of brain development ^3^. Notably, homozygous deletion of *Cul3* in mice is embryonically lethal suggesting its crucial role during early development ^4^.

Interestingly, one of Cul3 adapter proteins, KCTD13, is located within a large 16p11.2 CNV that confers high risk for neurodevelopmental disorders ^5^. We previously demonstrated that *Cul3* is co-expressed with *KCTD13*, and corresponding proteins physically interact during late mid-fetal period of brain development ^6^. This interaction is crucial for regulating the levels of a small GTPase RhoA that controls actin cytoskeleton structure and cell movement ^7^. Upregulation of RhoA through its constitutive expression has been linked to suppression of dendritic spine morphogenesis and to dramatic loss of spines ^8, 9^. The contribution of RhoA to local regulation of axon growth has been recently demonstrated ^10^.

To investigate how *Cul3* mutations impact early brain development at the molecular level, we generated *Cul3* haploinsufficient (*Cul3^+/-^*) mouse model using CRISPR/Cas9 genome engineering. We introduced a germline 1bp insertion in exon 6 of *Cul3,* immediately adjacent to G754T (E246X) mutation detected in an ASD patient ^11^ (**Figure 1b**). We extensively characterized *Cul3^+/-^* mice at multiple levels including transcriptomic and proteomic profiling of various brain regions at three developmental time points. Our results demonstrate that *Cul3* haploinsufficiency severely impacts early brain development through dysregulation of neuronal actin and intermediate filament cytoskeleton. Reduced dendrite growth and network activity of cortical neurons were among most notable neuronal phenotypes observed in *Cul3^+/-^* mice. Importantly, upregulation of one of the Cul3 substrates, small GTPase RhoA, a regulator of cytoskeletal dynamics, neuronal growth and migration, linked observed molecular defects with Cul3 pathology. Treatment of cortical neurons with RhoA inhibitor Rhosin rescued dendritic length and neural network activity phenotypes.

## Results

### *Cul3^+/-^* mice have brain anatomical defects starting from early postnatal period

To investigate the impact of *Cul3* mutations on brain development at the molecular level, we generated *Cul3^+/-^* haploinsufficient mouse model on C57BL/6N background by introducing a 1bp frame-shifting insertion into exon-6 of *Cul3* one base pair upstream from the nucleotide mutated in an ASD patient ^11^ using CRISPR/Cas9 genome editing (**Materials and Methods**). We validated the presence of mutation by Sanger sequencing (**Figure 1b**). In agreement with previous observations from conditional *Cul3^flox^* mice ^4^, homozygous *Cul3^−/−^* mutant animals were not viable. A heterozygous *Cul3^+/-^* founder mouse harboring the insertion was expanded via breeding with wild-type (WT) C57BL/6N mice for at least four generations to eliminate possible CRISPR off-target effects. We verified reduced levels of *Cul3* transcripts in F_4_ *Cul3^+/-^* mice by qRT-PCR (**Figure 1c**), and reduced Cul3 protein levels by Western blot (**Figure 1d**). These experiments demonstrated the reduction of Cul3 expression in the brain of *Cul3^+/-^* mutant mice to approximately half of the level observed in WT littermates, thereby validating *Cul3* haploinsufficiency.

The heterozygous *Cul3^+/-^* mice were viable, reached a normal lifespan, and were fertile irrespective of sex. However, *Cul3^+/-^* embryos (at both E15.5 and E17.5) and adult mice were smaller (**Figure 1e, Supplementary Fig. S1a**). Adult mutant mice had slightly visually elongated brain and significantly reduced body weight (P<0.001 for male and P<0.01 for female; two tailed t-test, **Figure 1f**).

To investigate in more detail whether Cul3 mutation impacts brain neuroanatomy, we analyzed PFA-fixed 8- to 10-week old adult WT and *Cul3^+/-^* mouse brains (N=38) using magnetic resonance imaging (MRI) (**Method Details**). We observed profound brain volume abnormalities in 46% (83/182, FDR<5%) of brain regions in *Cul3^+/-^* adult mice (**Figure 2a, Supplementary Table S2**). Total brain volume, as well as relative grey matter volume, were both significantly lower in *Cul3^+/-^* mice compared to WT (P<0.01 and P<0.05, respectively, two tailed t-test, **Figures 2b-c**). No significant differences in the white matter volume were observed between *Cul3^+/-^* and WT mice, with a trend for lower volume in *Cul3^+/-^* mice (**Figure 2d**). With respect to specific brain regions, several areas of somatosensory cortex, hippocampal dentate gyrus and cerebellum were significantly decreased in *Cul3^+/-^* mice (**Figure 2e**). Among other notable brain areas, many olfactory bulb regions, primary visual cortex, entorhinal cortex and corpus callosum were also decreased (**Supplementary Table S2).** Overall, the volume of many cortical regions was decreased, whereas the volume of some subcortical regions was increased in the adult mutant mice.

**Figure 2.**
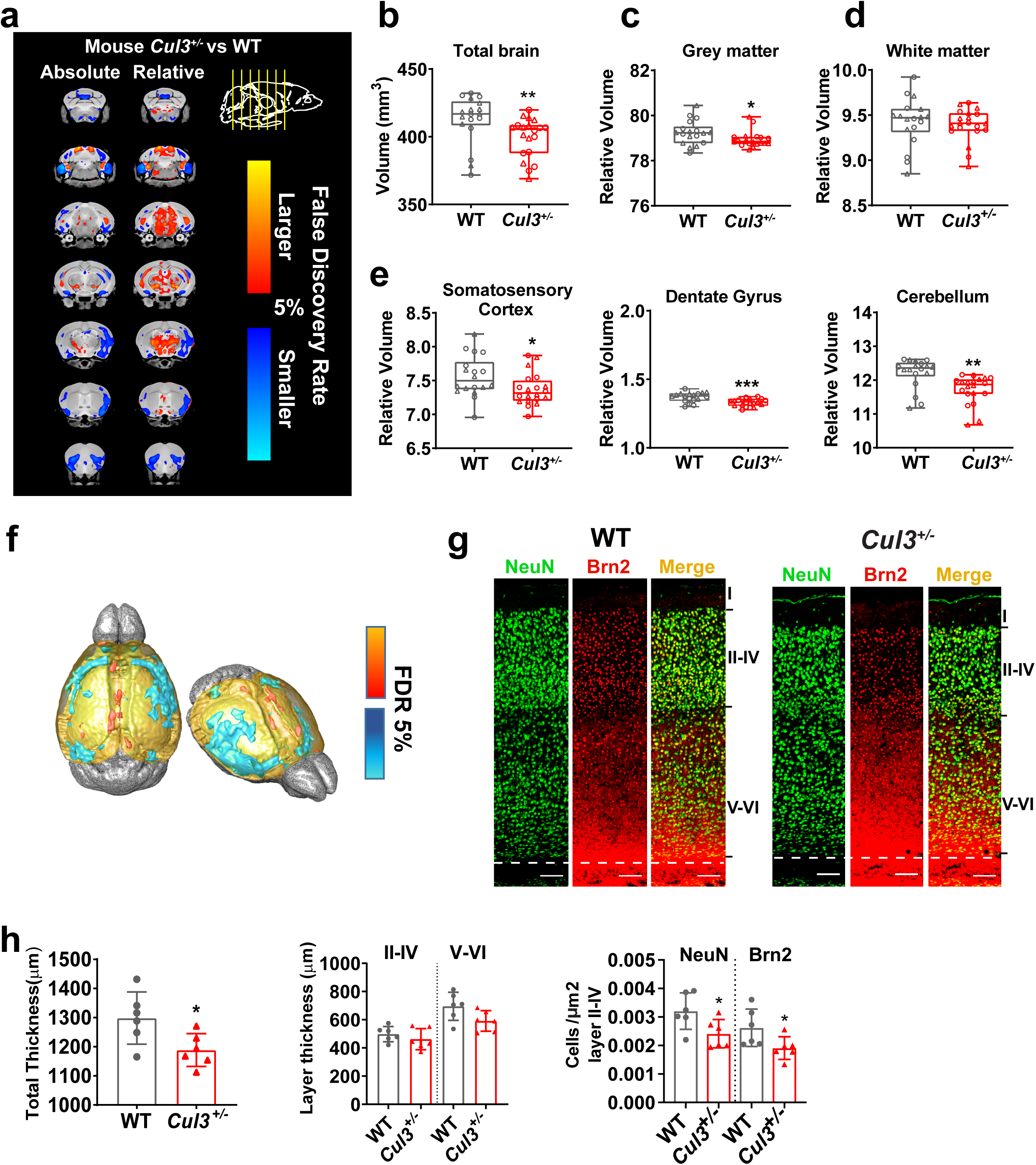
*Cul3^+/-^* mice have altered brain morphology. **(a)** Voxel-wise analysis highlighting significant differences in relative volumes throughout the brain between WT and *Cul3^+/-^* mice with 5% false discovery rate (FDR). Left panel is absolute changes and right panel is relative change in brain regions. Scale bar 2.7-10.7 indicates decreasing FDR, where 2.7=5% FDR. Red color signifies increased and blue color signifies decreased brain volume compared with WT brain. **(b)** *Cul3^+/-^* mice have reduced absolute brain volume (**p<0.01) compared to WT mice (WT n=18, *Cul3^+/-^ n*=20). **(c)** Reduced relative gray matter volume in *Cul3^+/-^* mouse brain (*p<0.05). **(d)** No significance difference is observed in relative white matter volume. **(e)** MRI revealed significant reduction in relative volume, normalized by absolute volume, of primary somatosensory cortex (*p<0.05); hippocampal DG region (***p<0.001); cerebellum (**p<0.01). Dots represent individual animals; ○-female and Δ-male**. (f)** Reduced cortical thickness is observed by MRI in *Cul3^+/-^* mice, turquoise color is decreased, red/orange color is increased. **(g)** Reduced cortical thickness of somatosensory cortex in *Cul3^+/-^* mice, left panel shows somatosensory cortex region stained with NeuN (mature neuron marker) and BRN2 (layer II-IV marker); Scale bar is 100μm. (**h**) Reduction of total cortical thickness, layer thickness and density of NeuN- and BRN2-positive cells/area in layer II-IV are observed (*p<0.05; two tailed T-test, n=6 for each genotype). Dots represent independent samples; two tailed T-test used for **b-h**; error bars represent mean ± SD.

To understand whether brain changes in mutant mice initiate early in development, we also performed MRI on postnatal day 7 (P7) fixed WT and *Cul3^+/-^* mouse brains (N=41) (**Supplementary Fig. S1, Supplementary Table S2**). Similar trends in brain volume changes were found in early postnatal P7 brain, with significant decrease in total brain volume, frontal cortex, parieto-temporal cortex, and cerebellum in mutant mice. Hypothalamus was increased in P7 mice. These results suggest that the impact of *Cul3* mutation initiates during early brain development.

The MRI data suggests that besides volumetric changes, *Cul3^+/-^* mice displayed decreased cortical thickness, particularly within frontal cortical regions of the brain (**Figure 2f**). To investigate what layers contributed to the observed reduced thickness, we immunostained brain sections with mature neuron marker NeuN and upper layer neuron (II-IV) marker BRN2 (**Figure 2g**). We observed significant overall reduction of cortical thickness in mutant mice (P<0.05, two tailed t-test, **Figure 2h,** left panel), even though the thickness of individual layers (II-IV and V-VI) demonstrated the trend but did not reach statistical significance. In addition, we observed reduction in NeuN and BRN2 positive neuron number in layers II-IV in the mutant mice (P<0.05, two tailed t-test, **Figure 2h,** right panel). This reduction in neuron number is likely due to premature neuronal death as evidenced by increased apoptosis measured by terminal deoxynucleotidyl transferase dUTP nick end labeling (TUNEL) in the primary cortical neurons at DIV14 (**Supplementary Figure S2**). In summary, Cul3 mutation impacts overall embryonic development along with brain volume, cortical thickness and neuron number starting from early postnatal periods.

### *Cul3* mutation impacts the behavior of Cul3^+/-^ mice

Although the main goal of this study was to identify molecular pathways disrupted by Cul3 mutation, we performed limited behavior testing of *Cul3^+/-^* animals for hyperactivity and social interaction. We would like to emphasize that complete behavior characterization of this mouse models was not the goal of this study. In the open field test, *Cul3^+/-^* mice travelled longer distances (P=0.032; two tailed t-test, **Figure 3a**) with higher speed (P=0.032; t-test; **Figure 3b**), suggesting hyperactivity. Data analysis in 10 min time bins revealed that hyperactivity of mutant animals increased after 10 min from the beginning of the test (**Figure 3c**). To assess whether hyperactivity is related to anxiety, we measured time spent in the center of the open field arena *vs* periphery, and observed no difference between WT and *Cul3^+/-^* animals (**Supplementary Figure S3**). Self-grooming that may also suggest anxiety was not increased in the mutant mice (**Figure 3d**). These results suggest that *Cul3^+/-^* mice are hyperactive but not anxious.

**Figure 3.**
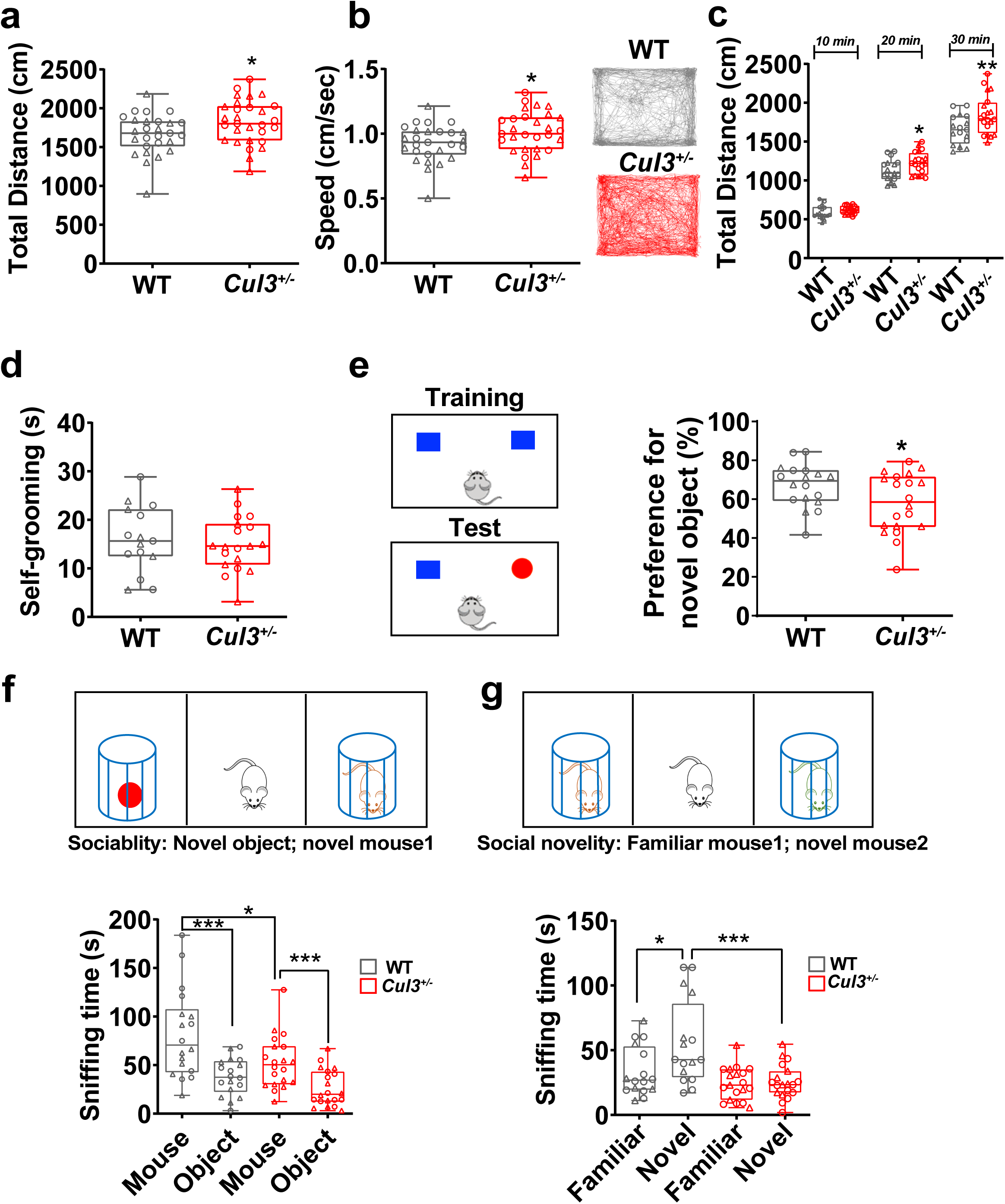
*Cul3^+/-^* mice display hyperactivity, cognitive and social impairments. **(a)** *Cul3^+/-^* mice travel longer distances in open field (n=27 WT; n=29 *Cul3^+/-^* (*p<0.05)) compared with WT mice. **(b)** The travelling speed (cm/sec) is significantly increased in *Cul3^+/-^* mice (*p<0.05); representative traces of 30 min in open field show *Cul3^+/-^* mice traveling longer distances compared to WT mice. **(c)** Time bins showing that *Cul3^+/-^* mice travel significantly longer distance in the last 20 min of the test (* p<0.05; **p<0.01). **(d)** No difference in time spent self-grooming between WT and *Cul3^+/-^* mice (WT n=15; *Cul3^+/-^* n=19). **(e)** *Cul3^+/-^* mice demonstrate significantly reduced preference for novel object in novel object recognition test (*p<0.05; Two tailed T-test, WT n=18; *Cul3^+/-^* n=20). **(f)** (Upper panel) Schematic diagram of three chamber social interaction test; reduced sniffing time is observed for *Cul3^+/-^* mice while interacting with a novel mouse (*vs* novel object) as compared to WT mice (WT n=17; *Cul3^+/-^* n=19; *p<0.05; One-way ANOVA). **(g)** Reduced sniffing time is observed for *Cul3^+/-^* mice while interacting with a novel mouse (*vs* familiar mouse) as compared to WT mice (WT n=17; *Cul3^+/-^* n=19) ***p<0.01; One-way ANOVA. Dots represent individual animals; ○-female and Δ-male; error bars represent mean ± SD.

Autism spectrum disorder is characterized by impairment in multiple domains including memory and social functioning. To investigate whether Cul3 mutation impacts learning, short-term memory, and social interaction, we used novel object recognition test. In this task, *Cul3^+/-^* mice spent shorter time exploring novel object than WT mice (P<0.05, two tailed t-test, **Figure 3e**). We then performed a three-chamber social interaction test, in which we first measured the sociability by analyzing the sniffing time of novel mouse *vs* novel object. Both *Cul3^+/-^* and WT mice preferred the mouse to the object, but *Cul3^+/-^* mice spent slightly less time sniffing the mouse than WT mice (P<0.001, one way ANOVA, **Figure 3f**). However, when a novel mouse was introduced instead of the object in the social novelty phase, *Cul3^+/-^* mice spent the same time sniffing novel and familiar mouse, and this sniffing time was shorter in mutant compared to WT mice (P<0.001, one way ANOVA, **Figure 3g**). These results suggest that *Cul3^+/-^* mice have impaired short-term memory and social interaction.

Since we observed changes in brain anatomy starting from early postnatal period, we investigated general developmental milestones in newborn mice through 21 days of age, in addition to behavioral testing of adult mice. There were no differences between genotypes in the eye opening, ear-twitching, auditory startle, forelimbs grasping and other developmental milestones (**Supplementary Figure S4**). The *Cul3^+/-^* mice demonstrated delayed performance in surface righting and cliff aversion tests, suggesting potential problems with motor control and vestibular difficulties. However, motor control deficiency did not persist into adulthood as demonstrated by longer distance traveled and higher speed in the open field test for adult mutant mice (**Figure 3a-c**).

### *Cul3* mutation dysregulates neurogenesis and cytoskeletal genes transcriptome-wide

Prior to investigating the impact of *Cul3* mutation on the developing mouse brain, we first analyzed *Cul3* expression in publicly available human and mouse brain transcriptome datasets. In human, *Cul3* gene is highly expressed in neocortex during early embryonic development (8pcw-13pcw), gradually decreasing in late fetal and especially postnatal periods according to BrainSpan, the transcriptome of the developing human brain ^3^ (**Supplementary Figure S5**). In mouse, *Cul3* expression was also higher in embryonic periods than in postnatal or adult, suggesting that *Cul3* gene may play important role in early brain development. Single cell RNA-seq profiling of human and mouse brain demonstrated that Cul3 is expressed at higher levels in neurons compared to other cell types in both species, with second highest expression in microglia/microphage in human, but not in mouse. This suggests that disruption of *Cul3* expression may have greater impact on neuronal cell types.

Since *Cul3* expression is the highest at embryonic periods, and we have observed abnormalities in both, early postnatal and adult brain, we performed bulk RNA sequencing (RNA-seq) of 108 brain transcriptomes derived from three developmental periods (embryonic E17.5, early postnatal P7 and adult 4-6 weeks) and three brain regions (cortex-CX, hippocampus-HIP and cerebellum-CB) of *Cul3^+/-^* mutant and WT mice (**Supplementary Figure S6**). The goal of transcriptome profiling was to detect how *Cul3* mutation impacts other genes at the transcriptional level. All three selected brain regions demonstrated decreased volume on MRI, and were previously implicated in ASD. After rigorous quality control of the data (**Supplementary Figure S7**), we performed differential gene expression analyses to identify genes that were up- or down-regulated in the *Cul3^+/-^* mutant *vs* WT mice (**Method Details**).

As expected, *Cul3* was the most differentially expressed protein-coding gene in all datasets, validating haploinsufficiency **(Figure 4a-c and Supplementary Figure S8)**. Dosage changes of *Cul3* gene also had transcriptome-wide *trans*-effect, dysregulating other genes. We identified hundreds of differentially expressed genes (DEG) between *Cul3^+/-^* mutant and WT mice at 10% FDR across developmental periods and brain regions (**Supplementary Table S3**). We identified 736, 1239 and 1350 unique DEGs in embryonic, early postnatal and adult periods, respectively. Likewise, we identified 727, 1641 and 1032 unique DEGs in CX, CB and HIP, respectively. Higher number of shared DEGs was observed between embryonic *vs* early postnatal (N=226) and early postnatal *vs* adult (N=255) periods, with the lowest number in embryonic vs adult (N=156) **(Supplementary Figure S8)**. Likewise, higher number of shared DEGs was observed between CB *vs* HIP (N=362), followed by CX *vs* CB (N=182) and CX *vs* HIP (N=159). Gene Ontology (GO) functional annotations of DEGs revealed enrichment of terms shared among multiple periods and regions as well as those that are unique to specific periods and regions (**Supplementary Table S4**). For example, GO functions of DEGs in CX included mostly neuronal processes (neuron projection, dendritic spine morphogenesis, axonogenesis), whereas HIP DEGs were enriched in cilium and cell adhesion. On the other hand, the DEGs in CB were enriched in synaptic and ion transport GO functions.

**Figure 4.**
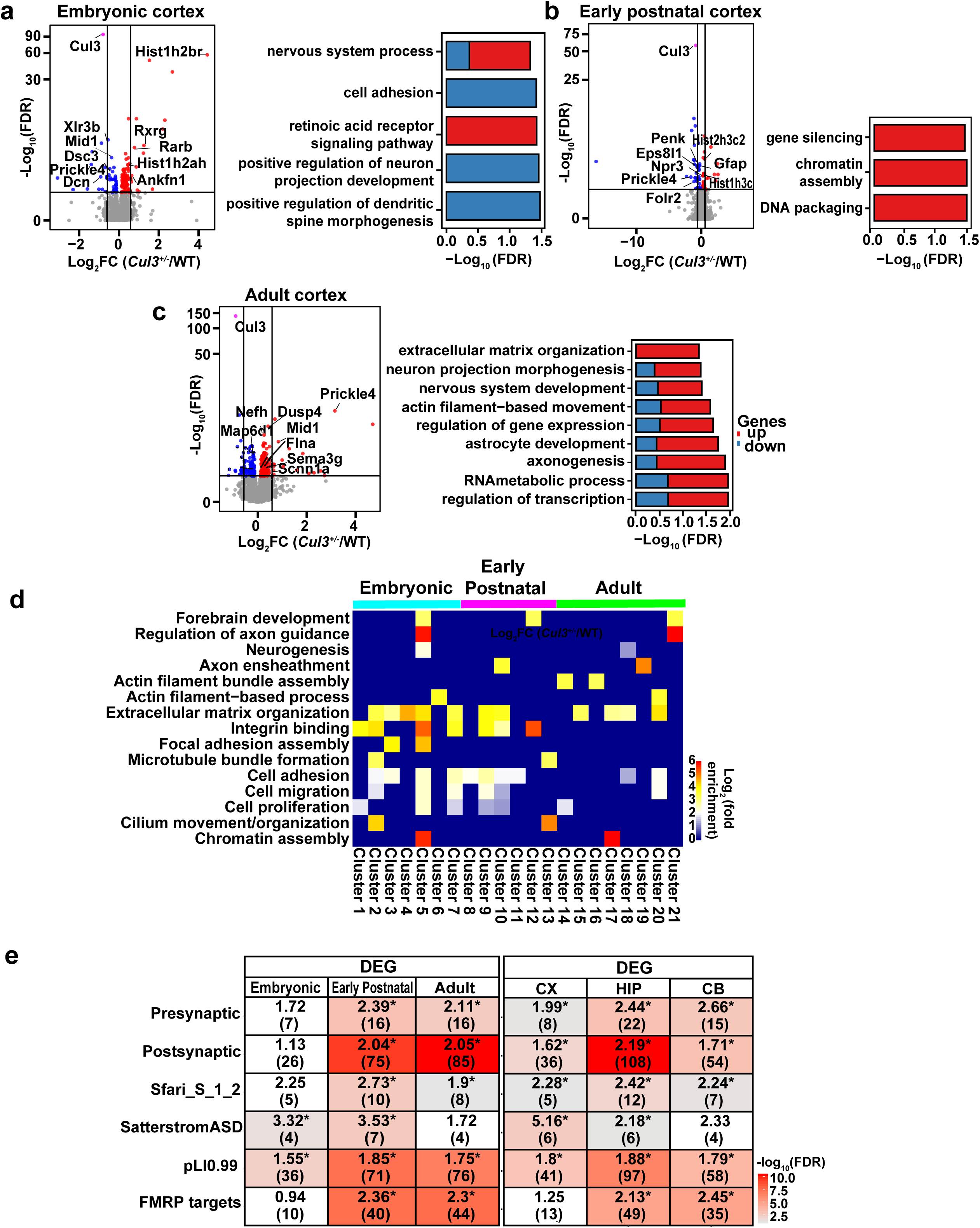
Spatiotemporal differential gene expression analyses identifies dysregulation of cytoskeletal processes by *Cul3* mutation. **(a)** Differential gene expression analyses of cortical samples from embryonic, early postnatal and adult developmental periods. (Left) Volcano plots of differentially expressed genes in *Cul3^+/-^ vs* WT. Genes colored in red are upregulated in *Cul3^+/-^* compared to WT; genes colored in blue are downregulated in *Cul3^+/-^* compared to WT; Cul3 is colored in pink. (Right) GO-terms enrichment of differentially expressed genes. Contribution of up- or down-regulated genes to specific GO terms are shown in blue and red, respectively. **(b)** Heatmap of enriched common GO terms at different developmental time periods. Fisher P-value combination analysis **(Materials and Methods**) was applied to identify 21 clusters of differentially expressed up- and down-regulated genes impacted by Cul3 mutation across developmental periods. Biological processes impacted in two or more clusters are shown (Individual cluster details are present in **Supplementary Table S6**). **(c)** Enrichment of differentially expressed genes from each period and region with literature-curated gene lists with previous evidence for involvement in autism. These lists include pre- and post-synaptic genes from SynaptomeDB; syndromic and highly ranked (1 and 2) genes from SFARI Gene database (https://gene.sfari.org/database/gene-scoring/); genes with probability of loss-of-function intolerance (pLI)>0.99 as reported by the Exome Aggregation Consortium; constrained genes; and FMRP target genes. Number of overlapped genes (in parenthesis) and odds ratio are indicated inside each cell, and provided only for FDR<0.05 and OR>1.

To better understand how gene expression is disrupted by *Cul3* mutation across all periods and regions, we performed meta-analyses of DEGs (**Materials and Methods**). This combined analysis identified clusters of genes that were either up- or down regulated in all periods or regions (**Supplementary Table S5**). GO annotation of these clusters highlighted terms shared by at least two clusters (**Figure 4d and Supplementary Table S6**). Biological process such as neurogenesis and axon guidance, integrin and extracellular matrix, cell adhesion, migration and proliferation, as well as actin cytoskeleton were altered in multiple clusters suggesting that Cul3 affects these processes transcriptome-wide.

To put our DEGs into a context of the existing knowledge of ASD genetics, we performed statistical enrichment analyses of DEGs against curated gene lists with previous evidence for involvement in ASD (**Figure 4e**). We observed enrichment of embryonic and early postnatal DEGs in high confident ASD risk genes from Satterstrom (OR>3) ^12^. Early postnatal and adult DEGs were enriched in presynaptic, postsynaptic and FMRP binding targets. DEGs from all three periods were enriched in genes highly intolerant to mutations (pLI>0.99). With regards to regions, CX was enriched in ASD risk genes from Satterstrom (OR>5), and all three brain regions were enriched in presynaptic and postsynaptic, ASD risk genes from SFARI, and genes highly intolerant to mutations (pLI>0.99). This suggests that *Cul3* mutation dysregulates genes relevant to ASD pathogenesis, especially in cortex and during embryonic and early postnatal periods. This is consistent with human data highlighting the impact of ASD mutations during late mid-fetal cortical development ^13, 14^.

### Proteomic profiling supports neuron cytoskeleton and neuron projection development dysregulation

Cul3 is a part of the ubiquitin-proteasome system that has significant role in protein turnover. Cul3 ubiquitin ligase interacts with a number of adapter proteins to ubiquitinate various substrates and to direct them to proteasomal degradation. Given the role of Cul3 in posttranscriptional regulation, we expect that mutations in Cul3 could impact cell’s proteome to a greater degree than they impact the transcriptome. To investigate the impact of Cul3 on brain proteome, we carried out quantitative Tandem Mass Tag mass spectrometry (TMT-MS) on 48 brain samples derived from three developmental periods (embryonic E17.5, early postnatal P7 and adult 4-6 weeks) and two brain regions (cortex CX and cerebellum CB) of *Cul3^+/-^* mutant and WT mice (**Supplementary Figure S9**). Differential protein expression analyses identified hundreds to thousands of differentially expressed proteins (DEPs) across various datasets (**Supplementary Table S7**). Overall, a greater number of DEPs was detected in embryonic brain *vs* early postnatal or adult, and in CX *vs* CB (**Supplementary Figure S10**).

GO annotations pointed to dysregulation of neuron projection, synaptic signaling and cytoskeletal functions **(Figure 5a)**. The neuron projection, intermediate filament and ion transport GO functions were shared among different periods (**Supplementary Table S8**). Notably, three cytoskeletal proteins, plastin 3 (Pls3), and neuronal intermediate filament proteins internexin (Ina) and vimentin (Vim), were found to be upregulated in all proteomics datasets, supporting neuron cytoskeleton dysregulation by Cul3 mutation with higher confidence (**Figure 5b**). We confirmed Pls3 upregulation by Western blot (**Supplementary Figure S10**).

**Figure 5.**
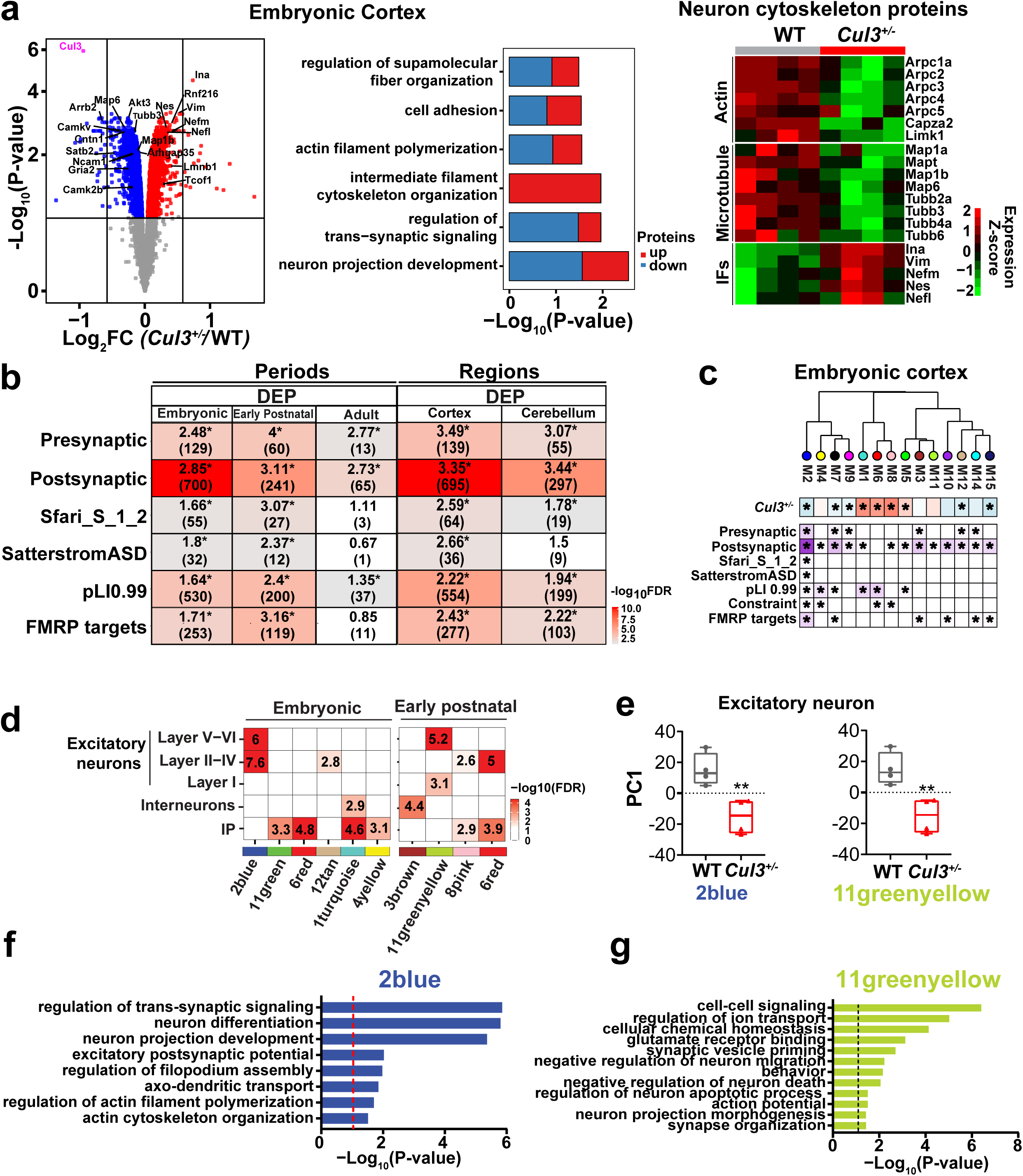
Differential protein expression and weighted protein co-expression network analyses of *Cul3^+/-^* mice. **(a)** (Left panel) Volcano plot of differentially expressed proteins between *Cul3^+/-^* and WT embryonic cortex identified from quantitative TMT-MS proteomic profiling. Cul3 is colored in pink, proteins colored in red are upregulated and proteins colored in blue are downregulated in *Cul3^+/-^*embryonic cortex. (Middle panel) Gene Ontology enrichment analyses of up- and downregulated proteins are shown as bar plots. Contribution of up- or down-regulated proteins to specific GO terms are shown in blue and red, respectively. (Right panel) Expression Heatmap of proteins associated with actin, microtubule and intermediate filament cytoskeleton in WT and *Cul3^+/-^* embryonic cortex. **(b)** Enrichment of differentially expressed proteins (combined by period and region) with literature-curated gene lists with previous evidence for involvement in autism. Number of overlapped proteins (in parenthesis) and odds ratio are shown. **(c)** Hierarchical clustering of protein co-expression modules by module eigengene for *Cul3^+/-^* embryonic cortex. Module-genotype associations (* FDR<0.1) for each module are shown below dendrogram. A total of 9 modules were significantly associated with *Cul3^+/-^* genotype in embryonic cortex. Module enrichment analyses against literature-curated gene lists with previous evidence for involvement in autism are shown at the bottom (* FDR<0.05) inside each cell, and provided only for FDR<0.05 and OR>1. **(d)** Cell type enrichment of co-expression modules from embryonic cortex using P0 mouse cortex scRNA-seq dataset ^16^. Modules significantly enriched in at least one cell type are shown for embryonic and early postnatal time periods. (**e**) PC1 of representative modules with significant cell type enrichment plotted by genotype. Embryonic module *2blue* and early postnatal module *11greenyellow* from Cul3 are depleted in excitatory neurons. All comparisons between WT and *Cul3^+/-^* are significant using T-test statistics. (**f-g**). GO terms for *2blue* and *11greenyellow* modules were obtained using g:Profiler.

Among differentially expressed cytoskeletal proteins, only intermediate neurofilament proteins (Vim, Nes, Ina, Nefl, Nefm) were significantly upregulated, whereas neuron microtubule or actin filaments proteins (Tubb2, Tubb3, Map1b, Actn1, Actn4, Limk, Arpc3, Arpc4, Capza2) were downregulated in the mutant embryonic mouse cortex (**Figure 5a**). The fine balance of Intermediate neurofilaments, microtubules and actin cytoskeleton is required for early cell positioning, polarization and neuritogenesis ^15^. Dysregulation of these proteins suggests potential impact on neuron growth and development during early embryonic development of *Cul3^+/-^* cortex. Interestingly, small GTPase RhoA, one of the substrate protein for Cul3 and regulator of actin cytoskeleton and neurite growth, is also upregulated in embryonic cortex and cerebellum. In addition, several other proteins with important neuronal and developmental functions were shared among datasets. They include receptor for netrin Dcc, required for axon guidance, contactin-associated protein-like Cntnap2, and treacle ribosome biogenesis factor Tcof1, involved in embryonic development of the craniofacial complex. Similar to DEGs, we performed enrichment analyses of DEPs with genes previously implicated in brain development and ASD. Embryonic and early postnatal developmental periods demonstrated stronger enrichment of genes from essentially all tested datasets, including ASD risk genes (**Figure 5b**). Likewise, CX had stronger enrichments across all datasets than CB. This suggests that Cul3 mutation may have greater impact on proteins involved in early cortical growth and development.

Besides differentially expressed proteins, the modules of proteins with correlated expression (*i.e.* co-expression modules) could provide further insight into pathways disrupted by Cul3 mutation. The Weighted Protein Co-expression Network Analysis (WPCNA) identified multiple protein co-expression modules positively or negatively associated with *Cul3^+/-^* genotype (**Supplementary Table S9**). We detected twenty-one co-expression modules in CX (9 embryonic, 5 early postnatal and 7 adult) and eleven modules in CB (8 embryonic, 1 early postnatal and 2 adult) that were positively or negatively associated with *Cul3^+/-^* genotype (**Supplementary Table S9**). GO annotations of CX (**Supplementary Table S10**) and CB modules **(Supplementary Table S11**) were in agreement with biological functions identified from DEG and DEP analyses. For example, *8pink* module was upregulated in *Cul3^+/-^* embryonic CX and enriched in intermediate filament proteins (Vim, Ina, Nes). Likewise, downregulated *2blue* module was enriched in neuronal and synaptic functions (neuron projection development, neuron differentiation, regulation of trans-synaptic signaling), with many ASD risk genes found within this module (CTNNB1, DPYSL2, NRXN1, STX1B, TBR1). Several modules contained proteins associated with chromatin and histone modification functions (*1turquoise* and *6red* in embryonic CX, and *6red* in early postnatal CX), and these modules were all upregulated in the *Cul3^+/-^* mouse. Further analyses demonstrated significant enrichment of almost all modules in postsynaptic proteins (**Supplementary Figure S11**), whereas downregulated *2blue* module in embryonic CX was enriched in ASD risk genes from Satterstrom ^12^ (**Figure 5c**). In summary, the WPCNA analyses of Cul3 spatio-temporal brain proteome further confirm dysregulation of neuronal and cytoskeletal protein modules by Cul3 mutation.

We performed bulk RNA-seq in this study. However, single cell RNA-seq (scRNA-seq) data is available from the same strain of mice. To better understand what cell types are impacted by Cul3 mutation, we performed enrichment analyses of our CX modules with scRNA-seq data from the P0 developing mouse neocortex ^16^. We observed several modules enriched in various cell types, with strongest enrichment of excitatory neurons from cortical layers II-VI in embryonic CX *2blue* and early postnatal CX *11green-yellow* modules (**Figure 5d**). Both modules were downregulated in *Cul3^+/-^* mice (**Figure 5e**), and their GO functions consisted on neuronal and synaptic functions such as “neuron projection development”, “neuron differentiation” and “trans-synaptic signaling” (**Figure 5f-g**). These results suggest that Cul3 mutation may impact excitatory neurons in the developing cortex of *Cul3^+/-^* mice.

### *Cul3* haploinsufficiency leads to reduced dendritic length and decreased neuronal network activity in primary cortical neurons

The DEGs, DEPs and co-expression modules in our transcriptomic and proteomic analyses were consistently enriched in neurogenesis, neuron projection development, and synaptic GO functions. We observed up- or downregulation of a number of proteins involved in dendrite and axon growth (Map2b, Tubb3, EphA7, DPYSL2, RAB family proteins), as well as cytoskeletal proteins involved in regulation of actin polymerization from Arp2/3 and WASP complexes (ARPC2, ARPC3, EVL, BAIAP2). In addition, our data demonstrates that Cul3 dysregulates early brain development, with potentially greater impact on cortical neuron growth and development.

To investigate neuronal defects in *Cul3^+/-^* mice, we examined neuron morphology of primary cortical neurons derived from E17.5 embryonic *Cul3^+/-^* mutant and WT mice. We calculated total dendrite length and the number of neurites, along with soma size, of 14DIV neurons stained with Map2 dendritic marker (**Figure 6a**). Tracing of Map2-positive dendrites demonstrated reduction of total dendritic length and the number of neurites in *Cul3^+/-^* mice (P=0.001; two tailed t-test; **Figures 6b-c**). Soma size was not affected by the *Cul3* mutation (**Figure 6d**).

**Figure 6.**
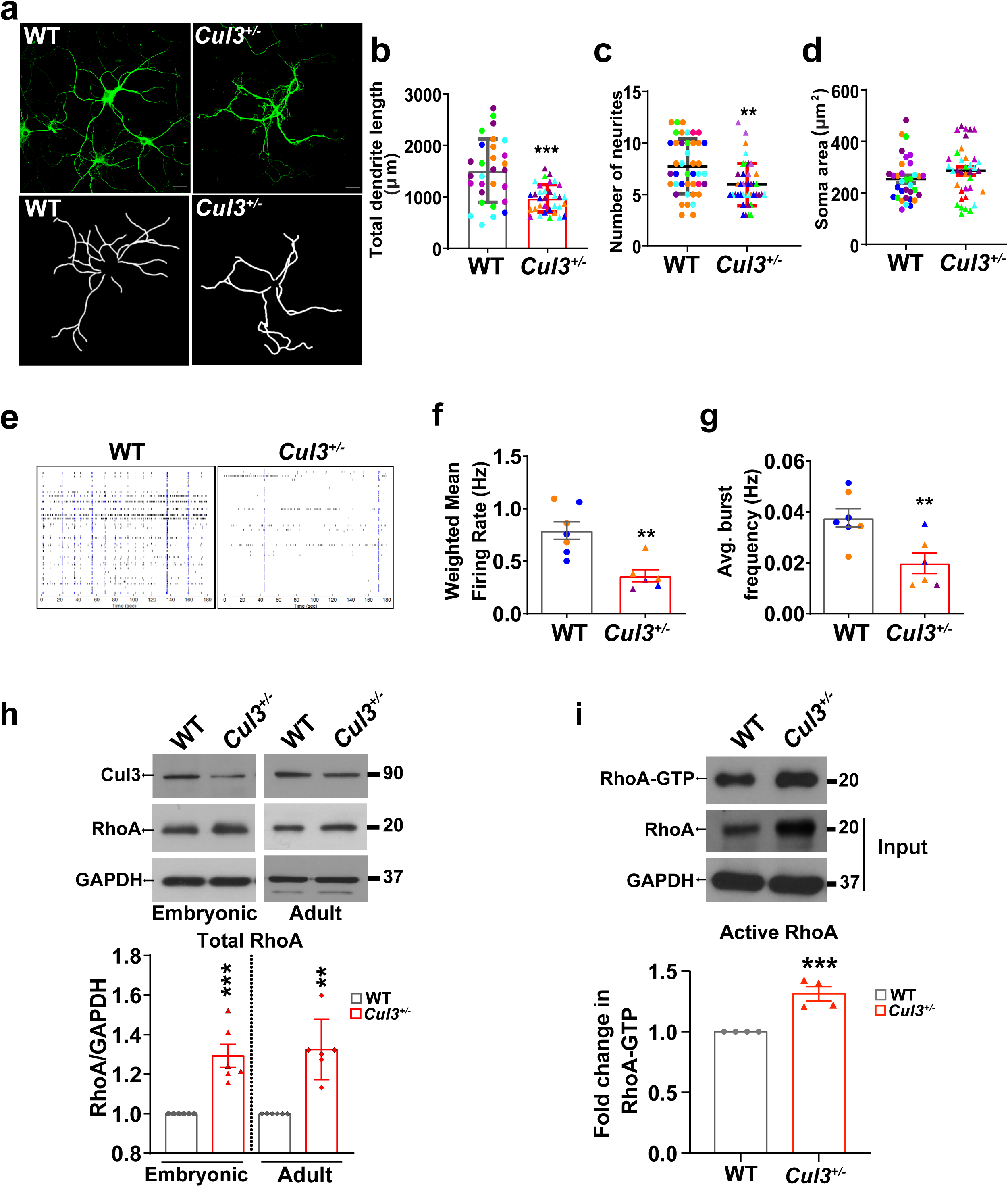
*Cul3* haploinsufficiency leads to altered neuron growth, network activity and to RhoA upregulation in *Cul3^+/-^* mice. **(a)** Representative images of 14DIV primary cortical neurons from WT and *Cul3^+/-^* mice, immunostained with MAP2 (upper panel), and tracings by simple neurite tracer (lower panel). Scale bar is 25 μm. **(b-d)** Quantification of total dendrite length, neurite number and soma size are shown. Symbols represent independent neurons and color represents littermates. Data is shown as mean ± SD (*n=*6 per genotype, at least 6-8 neurons per mouse). Significance is calculated using two tailed T-test; ***p<0.001; **p<0.01**. (e)** Representative raster plots of spontaneous spike activity from 8DIV primary cortical neurons. **(f)** Spontaneous spike activity is significantly reduced in *Cul3^+/-^* cortical neurons; **p <0.01, two tailed T-test. **(g)** Average burst frequency is significantly reduced in *Cul3^+/-^* neurons; **p< 0.01; two tailed T-test, n=6-7 mice per genotype; 1.5×10^6^ neurons were seeded from each mouse in each MEA plate well, each containing 64 electrodes. Each dot represents independent mouse and color represents littermates. **(h)** Representative images of Western Blot analysis of Cul3, total RhoA, and GAPDH loading control in embryonic and adult cortices. Densitometry analysis of Western Blot is shown at the bottom. Data is presented as mean ± SEM (n=6 per genotype). **(i)** Representative images of Western Blot analysis of active RhoA (RhoA-GTP) pulldown, total RhoA and GAPDH loading control from input lysate of embryonic cortex. Densitometry analysis of Western Blot is shown at the bottom. Data is presented as mean ± SEM (n=4 per genotype for total RhoA, n=4 per genotype for active RhoA). Significance is calculated using two tailed T-test; **p<0.01. Significance above bars represents comparison against WT.

Altered neuron morphology, and especially decreased dendritic length, could impact synaptic connectivity and alter neuronal network activity. We next examined network activity using multielectrode array (MEA) recordings in 8 days old primary cortical neuronal culture. Representative raster plots of spontaneous spike activity from these cultures are shown in **Figure 6e**. Analyses of spontaneous neural recordings demonstrated significantly reduced weighted mean firing rate in *Cul3*^+/-^ neurons compared to WT (P=0.002, two tailed t-test; **Figure 6f**). *Cul3*^+/-^ neurons also had significantly reduced bursting activity (P = 0.007, two tailed t-test; **Figure 6g**).

We next sought to search for the molecular target involved in dendritic deficits and reduced network activity of *Cul3^+/-^* cortical neurons. Based on our data, there are several indicators that point to potential dysregulation of Rho signaling in *Cul3*^+/-^ mouse brain. First, proteomic profiling identified a small GTPase RhoA as significantly upregulated protein in embryonic cortex and cerebellum. Second, RhoA is one of the direct molecular substrates of Cul3 ubiquitin ligase complex. Cul3 ubiquitinates RhoA and directs it for proteasomal degradation ^7^. Third, RhoA is an important regulator of actin cytoskeleton, neuron growth and migration during early brain development ^8^. Since we observed defects in neuronal growth, neurogenesis and neuron cytoskeleton, we tested for potential involvement of RhoA by Western blot in embryonic and adult CX. We observed significant upregulation of total RhoA in both tissues (P<0.001 for embryonic and P<0.01 for adult, two tailed t-test, **Figure 6h**). We also tested the level of GTP-bound active form of RhoA (RhoA-GTP) and observed its significant increase in embryonic CX, in agreement with total RhoA upregulation (P<0.001, two tailed t-test, **Figure 6i**). Increased RhoA levels during development may lead to growth cone retraction and altered neurite growth and extension.

### *Cul3* mutation destabilizes actin cytoskeleton and leads to loss of F-actin puncta in cortical neurons

Neuron cytoskeleton is a complex and elaborate structure, with actin, microtubules and intermediate filaments playing dynamic and coordinated roles in determining neuron shape, movement, and connectivity during embryonic development. The cytoskeleton is critical for formation of specialized structures such as growth cones, which are responsible for axon and dendrite elongation and guidance during development, synaptic boutons and dendritic spines, which form the structural basis for neural communication. Proteomic profiling of mouse brain pointed to neuronal cytoskeleton as one of the cellular structures impacted by *Cul3* haploinsufficiency. We found intermediate filament proteins to be upregulated across multiple DEP datasets in *Cul3^+/-^* mouse (**Figure 5a**), along with upregulated embryonic CX module *8pink* containing these proteins (**Figure 5c and Supplementary Table S10**). At the same time, proteins involved in actin cytoskeleton-related processes (actin filament binding, actin cytoskeleton organization, regulation of actin cytoskeleton polymerization, regulation of actin filament length) were found to be downregulated in *Cul3*^+/-^ brain. Specifically, we identified significantly downregulated *2blue* module in the embryonic and *7black* in the adult CX modules with actin cytoskeleton-related GO functions (**Figures 7a-b and Supplementary Table S10**).

**Figure 7.**
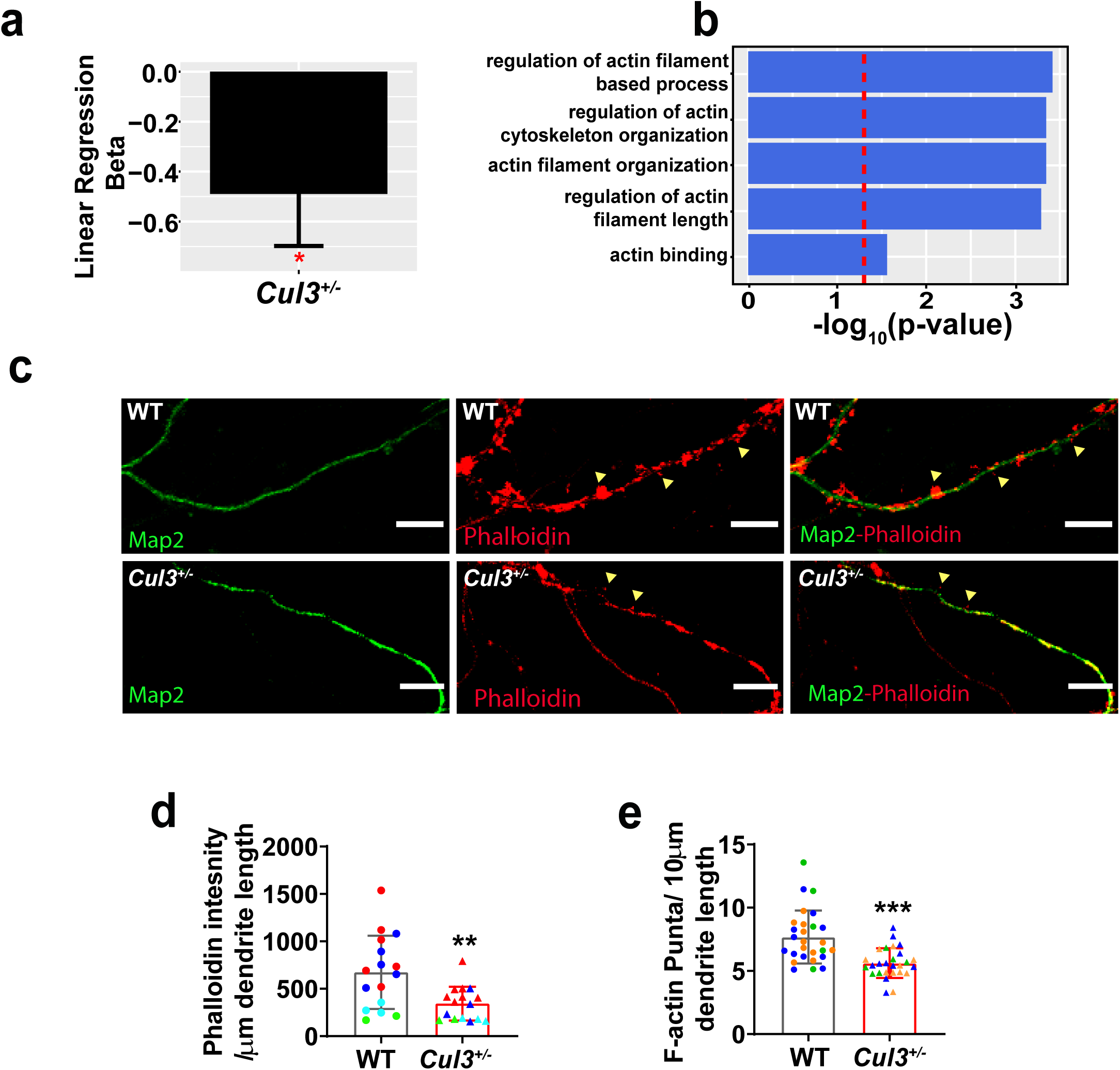
Actin cytoskeleton defects are observed in *Cul3^+/-^* mice. **(a-b)** Module-genotype association and GO functional annotations for *black* actin module identified by protein co-expression analyses of adult cortex. Dots represent independent animals (n=4 per genotype). **(c)** Representative images of 21DIV primary cortical neurons from WT and *Cul3^+/-^* mice, immunostained with MAP2 (green) and phalloidin-rhodamine (red); Scale bar is 25 μm; Δ represents F-actin puncta on dendritic segment for better visualization. **(d-e)** Quantification of F-actin puncta and F-actin intensity on MAP2 positive dendrites, normalized by dendrite length. Symbols represent independent neurons and color represents littermates. Data is presented as mean ± SD (n=4 per genotype, at least 7-10 neurons per mouse). Significance is calculated using two tailed T-test; ***p<0.001.

To validate potential actin cytoskeleton defects in *Cul3*^+/-^ mouse, we co-stained 21DIV cultured mature primary cortical neurons with Phalloidin-rhodamine and mature neuron marker Map2 to quantify filamentous actin (F-actin) (**Figure 7c)**. We observed significant reduction of Phalloidin intensity in *Cul3*^+/-^ mutant neurons (**Figure 7d**), as well as reduction of the number of F-actin puncta per dendrite length (**Figure 7e**). Dendrites shortening and filamentous actin loss likely impacted neural connectivity and lead to changes in neural network activity observed in the mutant mice by MEA.

### RhoA inhibition rescues dendritic growth and network activity of cortical neurons

Since RhoA regulates neuronal actin cytoskeleton remodeling and neurite outgrowth, and we found RhoA upregulated in *Cul3^+/-^* mice, we investigated whether RhoA inhibition could rescue decreased dendritic length and neural activity phenotypes in primary cortical neurons. We took advantage of a previously described pharmacological inhibitor of RhoA, Rhosin. Rhosin was shown to specifically block RhoA activation and induce neurite outgrowth ^17^. We repeated neuron morphometric analysis and used Rhosin (RH) or vehicle (VH) to treat primary cortical neuronal culture from 2 DIV until 14 DIV (**Figure 8a**). We replicated decreased dendritic length phenotype in VH-treated *Cul3*^+/-^ neurons, and Rhosin treatment rescued dendritic length defects in *Cul3*^+/-^ to the level indistinguishable from the WT (**Figure 8b-c**). Next, we measured network activity of primary cortical neurons at DIV8 with MEA in the presence and absence of Rhosin. We observed that Rhosin treatment was able to rescue the reduced weighted mean firing rate and burst activity, returning these parameters to the levels comparable with vehicle-treated WT cortical neurons (**Figure 8d-f**). Quality control included cell imaging to ensure comparable coverage of MEA plate, and protein quantification after completion of recording to ensure equal well coverage between genotypes (**Supplementary Figure S12**). These results suggest that Cul3 haploinsufficiency dysregulates neuron cytoskeleton and neurogenesis through increased active RhoA levels during cortical neuron development. The upregulation of active RhoA could be responsible for cellular, molecular and general neurodevelopmental defects observed in the *Cul3*^+/-^ mutant mice.

**Figure 8.**
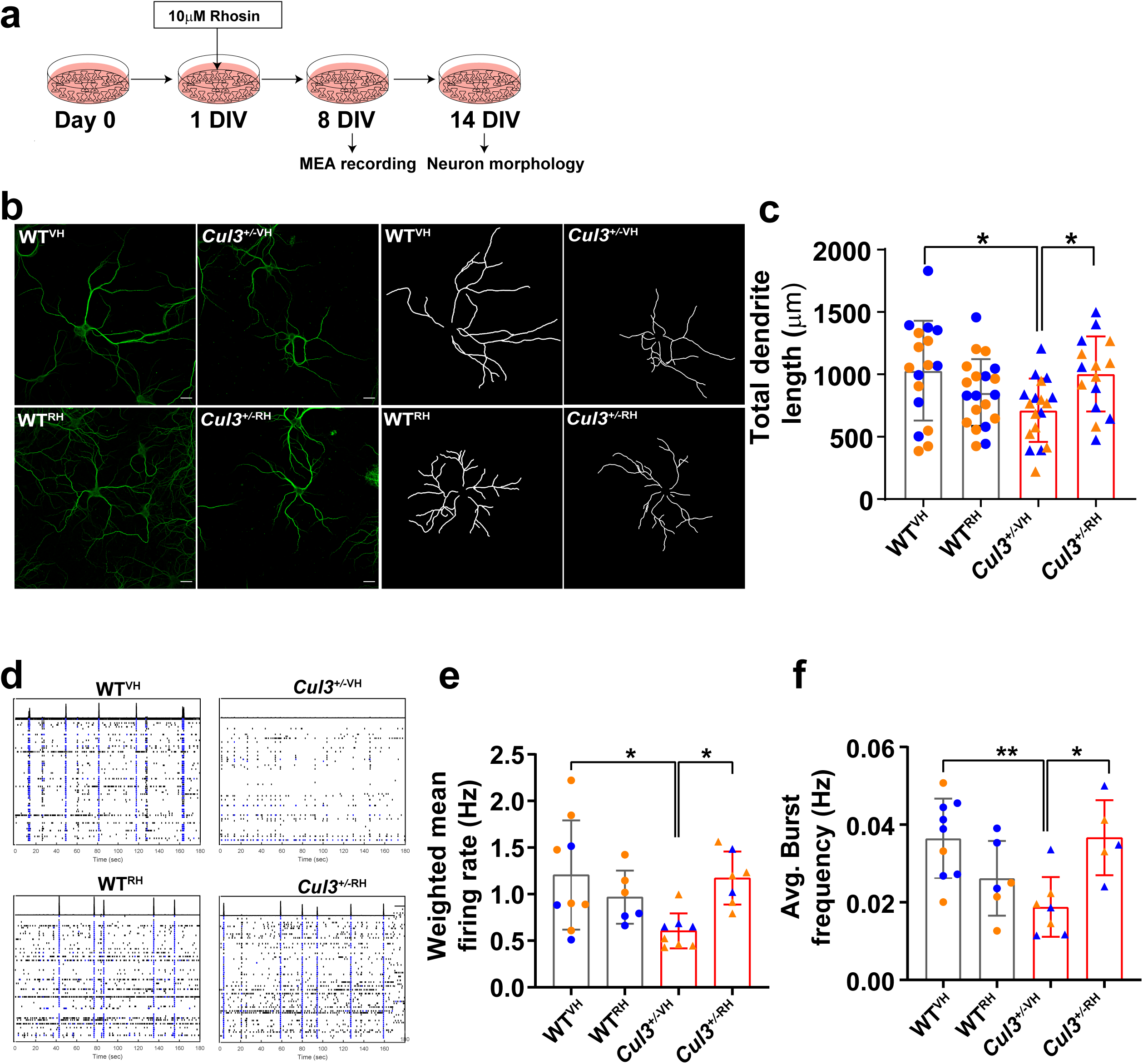
RhoA inhibition rescues dendritic growth and network deficits. **(a)** Flow diagram showing treatment timeline of primary cortical neurons with RhoA inhibitor Rhosin (RH). Half of the media containing Rhosin or vehicle was replaced every third day until the day of experiments. **(b)** Representative images of 14DIV primary cortical neurons (left panel) and tracings (right panel); Scale bar is 25 μm. The vehicle (Vh) and Rhosin (Rh) treated cells were immunostained with Map2 and dendrite tracing was performed. **(c)** Rhosin treatment rescues decreased dendrite length phenotype in *Cul3^+/-^* neurons. Symbols represent independent neurons and color represents littermates. Data is presented as mean ± SD (n=2 per genotype, at least 8-10 neurons per mouse). Significance is calculated using One way ANOVA and Tukey test for multiple comparison; *p<0.05. (**d**) Representative raster plots of spontaneous spike activity from 8DIV primary cortical neurons. **(e)** Spontaneous spike activity is significantly reduced in *Cul3^+/-^* cortical neurons and rescued by treatment with 10µM Rhosin; *p <0.05, One way ANOVA and Tukey test for multiple comparison. **(f)** Average burst frequency is significantly reduced in *Cul3^+/-^* neurons and rescued by treatment with 10µM Rhosin; n=6-9 mice per genotype, **p< 0.01; * p< 0.05. One-way ANOVA and Tukey test for multiple comparison was used.

## Discussion

Loss-of-function mutations in Cul3 ubiquitin ligase are strongly associated with autism and developmental delay. However, the molecular mechanism by which *Cul3* haploinsufficiency causes these diseases are unknown. Since *Cul3* is highly expressed in the embryonic human brain, and its expression decreases after birth, it is likely that its molecular impact is manifested during early brain development. In this study, we generated the first CRISPR/Cas9 mouse model carrying loss-of-function mutation in the same exon of *Cul3* gene as ASD patient’s mutation. We characterized this model using brain MRI, transcriptomic and proteomic profiling, and provide evidence implicating neuron cytoskeletal proteins and cortical neurogenesis along with RhoA signaling as potential mechanisms of *Cul3* haploinsufficiency.

At the neuroanatomical level, we observed changes in both early postnatal day 7 and adult brain of *Cul3*^+/-^ mutant mice, suggesting profound impact of *Cul3* mutation on the brain that begins early in development. The volume of many cortical structures (visual, somatosensory, hippocampus-CA2, dentate gyrus, corpus callosum and cerebellar cortex) was reduced in mutant animals. These changes in brain anatomy share similarities with other autism mouse models summarized in a previous study that clustered 26 models into three groups by correlated volume changes of distinct brain regions ^18^. *Cul3*^+/-^ model is most similar to the group comprised of 16p11.2, Mecp2, and CNTNAP2 mouse models. Comparable brain structural changes were later reported in a different 16p11.2 deletion mouse model ^19^, as well as in TAOK2 mouse model ^20^. These similarities between 16p11.2 and Cul3 models is not surprising because one of the genes from 16p11.2 CNV region, *KCTD13*, is an adaptor for Cul3 protein ligase. Thus, the same pathways may be dysregulated by the 16p11.2 CNV and Cul3 mutation.

In addition to brain volumetric changes, we observed decreased cortical thickness, neuron density and increased apoptosis of cortical neurons in the *Cul3*^+/-^ mouse. The scRNA-seq enrichment analysis demonstrated that the cortical excitatory neurons were most affected by Cul3 mutation. Neurons from upper layers are likely contributing to these phenotypes, given downregulation of proteins expressed in the upper layer neurons (Satb2 and beta-tubulin marker protein Tubb3). The upregulation of neuroprogenitor marker proteins (Sox2 and Nes) in the embryonic cortex of *Cul3*^+/-^ mutant mice points towards delayed maturation of *Cul3*^+/-^ mutant cortex. The reduced number of upper layer BRN2-positive neurons that is validated by the TUNNEL assay, suggests increased apoptosis. In addition, defects in neuronal migration, supported by our transcriptomic analyses, could also contribute to the decreased cortical thickness, although we have not investigated migration experimentally in this study.

Neuron morphology, and specifically dendrite length, was reduced in *Cul3*^+/-^ mice. Proteomic results identified module of co-expressed proteins with neuron projection development functions. The observed dendritic growth deficits of *Cul3*^+/-^ neurons likely influencing synaptic connectivity and network activity, as we observed reduced firing rate and bursting activity in these neurons. RhoA is one of the proteins that controls neurite outgrowth, and RhoA upregulation was previously implicated in loss of spines and neurite retraction ^6, 7, 10, 17, 21^. Previously developed genetic autism mouse models (SETD5, UBE3A, SHANK3, Timothy syndrome) also observed reduced neurite outgrowth, sometimes in concert with reduced synaptic connectivity ^21–24^. The iPSC-derived neurons from Timothy syndrome patients also exhibited dendrite retraction phenotype ^21^. Thus, reduced neurite outgrowth, possibly due to upregulated RhoA signaling, could be a shared phenotype among different autism models, including *Cul3*^+/-^.

Dysregulation of neuron cytoskeletal proteins in particular suggests a possible mechanism underlying *Cul3* haploinsufficiency. Neuronal cytoskeleton is crucial for a number of biological functions such as extracellular matrix organization, cell adhesion, microtubule formation, neuron migration, motility and locomotion, and cilium-related functions. We observed all of these processes to be enriched by either transcriptome or proteome profiling of Cul3^+/-^ brain. In addition, neuron cytoskeleton is essential for proper neurogenesis, branching and dendrite arborization, emphasizing the importance of cytoskeletal integrity during neocortical development ^25^. Cul3 impacted a number of cytoskeletal proteins, including those involved in intermediate filament, microtubule and actin cytoskeleton organization, remodeling, binding and regulation. Cytoskeletal proteins were altered in all developmental periods and regions, with three upregulated proteins (Vim, Pls3 and Ina) consistently shared by all datasets. Other proteins with actin-related functions formed a co-expression module downregulated in the mutant mouse. Published literature demonstrated that intermediate filaments play important role in promoting microtubule stabilization, and their accumulation results in hyperstabilization of microtubules, leading to disruption of synaptic vesicle transport ^26^. Vimentin is known to be involved in early neuron outgrowth and cell polarity. It also acts as a template for microtubule stabilization during cell migration ^27^. Kurup et al. proposed that increased neuronal apoptosis and defective neurite outgrowth may be the consequence of accumulated intermediate filaments that affect the transient actin and microtubule cytoskeleton. We experimentally confirmed loss of filamentous actin puncta in *Cul3*^+/-^ neurons, providing further evidence for cytoskeletal defects.

We attribute observed cytoskeletal and neuronal defects to upregulation of RhoA signaling. First, we observed RhoA upregulation in embryonic and adult cortex of *Cul3* mutant mice. Second, we were able to rescue dendrite length and network connectivity phenotypes by pharmacological inhibition of RhoA activity. Third, RhoA upregulation was found in other relevant ASD models, such as KCTD13 ^28^. KCTD13 is an adaptor for Cul3 that regulate RhoA ubiquitination and degradation. Thus, downregulation of either KCTD13 or Cul3 are expected to upregulate the levels of RhoA. A recent study of conditional *Cul3* model also observed RhoA upregulation in the cortex ^29^. Most importantly, RhoA is a key regulator of actin cytoskeleton remodeling, neurite outgrowth, migration and spine formation during early cortical development ^30^, and we observed defects in many of these pathways in *Cul3*^+/-^ mouse. It is likely that Cul3 also dysregulates other pathways upstream or downstream of RhoA to cause these and other phenotypes. This possibility could not be ruled out by our study and needs further investigation.

Finally, we compared our findings with two recently published Cul3 conditional mouse models ^29, 31^. These models were generated by either knocking down or knocking out Cul3 in excitatory neurons of prefrontal cortex ^29^, or by activating mutation at progenitor’s level, leading to complete elimination or haploinsufficiency in neurons and glia ^31^. The homozygous germline knockout Cul3 mice are embryonically lethal (^4^ and this study), therefore we resorted our comparison to heterozygous mice from these studies with our heterozygous *Cul3*^+/-^ model. Our Cul3 mouse model differs from two other models in at least two ways: (1) CRISPR technology allowed us to generate germline mutation that is present in all cell types and in all brain regions and other tissues, which is more comparable to human carriers; (2) Cul3 mutation is present starting from conception and from the very early developmental time points. Finally, none of two former models investigated the impact of Cul3 mutations in embryonic or early developmental periods, which is crucial due to highest expression of Cul3 gene during early prenatal development. The changes observed at later developmental time points in the published studies may be just the consequences of early embryonic deficits.

Interestingly, both studies reported reduced body weight only in Cul3 homozygous knockout mice, but not in heterozygous mutants, suggesting a milder phenotypes observed in the previous models compared to our CRISPR *Cul3*^+/-^ mouse. Dong model also observed hyperactivity, in agreement with our model. Previous models did not perform transcriptomic analyses, therefore we were able to only compare proteomics results from these models with our findings (**Supplementary Figure S13)**. Importantly, neither Dong nor Rapanelli studies detected differentially expressed proteins with FDR-corrected p-values in their proteomics data, because none of their proteins surpassed FDR correction. When genome- and proteome-wide studies are carried out, only genes/proteins with FDR-corrected p-values are normally reported to avoid spurious findings. We, on the other hand, have detected 2,329 differentially expressed proteins in embryonic, 986 in early postnatal and 402 in adult cortex; and 1,016 in embryonic, 252 in early postnatal and 5 in adult cerebellum with FDR<15%. This suggests that our study had greater power to detect differentially expressed proteins and to implicate molecular pathways, potentially due to stronger effect of Cul3 mutation due to its germline transmission. Clearly, the embryonic period stands out with much greater number of detected DEPs.

Importantly, Dong and Rapanelli studies did not converge on a common molecular mechanism underlying the impact of Cul3 mutation. Dong et al. observed translational dysregulation in their Cul3 model, with the main contributing player translation initiation factor EIF4G1. In our study, we did not observe EIF4G1 to be differentially expressed at any period or in any brain region, although it has been detected by proteomics. At the same time, our upregulated DEPs Pls3 and Ina were also upregulated in Dong model with nominal uncorrected p-values. Rapanelli et al. study narrowed down to histone methyltransferase Smyd3. We have not detected Smyd3 in our proteomics experiments, therefore we can not validate this specific finding. However, Rapanelli also observed RhoA upregulation and reduced spine density, in agreement with our results. Interestingly, Pls3 was the only protein that was found to be upregulated in Dong, Rapanelli and our study. We conclude that our study captures early molecular defects in cortical neuron cytoskeleton and neurogenesis that likely lead to molecular and physiological changes at later developmental time points observed by other models.

In conclusion, our study demonstrates that *Cul3* germline haploinsufficiency impairs early brain growth and development by dysregulating neuronal cytoskeleton and neural connectivity through a molecular mechanism related to RhoA signaling. Other pathways, detected later in development and in adulthood, are likely the result of these initial defects that begin during early embryogenesis.

## Acknowledgments

This work was supported by a grant to LMI and ARM from the National Institute of Mental Health (MH108528), and by grants to LMI and ARM (MH109885, Simons Foundation for Autism Research #345469), to LMI (MH105524, MH104766), to JRY and ARM (MH100175), and to JS (MH119746). RNA-seq data was generated at the UC San Diego IGM Genomics Center, University of California San Diego (grant P30CA023100). The images were acquired at the UCSD School of Medicine Microscopy Shared Facility (grant NS047101). We thank Dr. Joseph Gleeson and Dr. Lu Wang for help with Maestro instrument, Dr. Barun Das for TUNEL data analysis, Punit Mehta for help with mouse behavior data analyses, Dr. Alessandra Porcu and Dr. Ben Romoli for histology support, and the UCSD undergraduate students Kaiyi Mu, Jasmine Le, and Camille Sotelo for help with image analysis.

## Author contributions

L.M.I. and A.R.M. conceived the study. M.A., A.B.P., V.M.H., N-K.Y., L.R.Q., P.M-L., P.Z., C.A.T., J.E., J.U., K.C., J.D., J.S., D.R., J.P.L., J.R.Y. III, A.R.M. and L.M.I. designed the experiments and analyses. M.A., V.M.H., N-K.Y., L.R.Q., C.A.T., J.U., J.D., and J.G. performed experiments and analyses. A.B.P., P.M-L., P.Z., K.C., J.E., K.C., J.C., and D.R. performed computational data processing and analyses. M.A. and L.M.I. wrote the manuscript, with input from all co-authors. Supervision was performed by J.S., D.R., J.P.L., J.R.Y. III, A.R.M, and L.M.I.

## Lead contacts and material availability

Further information and requests for resources and reagents should be directed to and will be fulfilled by the Lead Contact, Lilia M. Iakoucheva. New *Cul3*^+/-^ mouse model will be shared with interested investigators through Jackson Laboratory as described in the NIMH Data Sharing agreement.

## Materials and Methods

### Generation of *Cul3^+/-^* mice using CRISPR-Cas9 genome editing and maintenance

For generating *Cul3* mouse model with the patient-specific p.Glu246Stop (E246X) mutation ^11^, single-guide RNAs (sgRNA) and single-stranded DNA oligonucleotides (ssODN) were designed to create the E246X frameshift mutation. Two single guide (Sg) RNA, Sg1-GGATTAATGAGGAAATAGAG and Sg2-AGTAGAAGCTAGGATTAATG, with little off-target specificity for *Cul3* (NM 016716.5) were designed using http://crispr.mit.edu/ platform and CHOP-CHOP (https://chopchop.cbu.uib.no/). The specificity and efficiency of each sgRNA was validated using *in vitro* Cas9 cleavage assay. Generation of transgenic lines was performed by the UCSD Moores Cancer Center Transgenic Mouse Core Facility. Briefly, microinjections consisting of sgRNA1 (0.6μM of crRNA, 0.6μM tracrRNA), 0.5μM of ssODN and 30ng/μl Cas9 protein (PNA-Bio, USA) was injected in pronuclear stage embryos in the C57BL/6N genetic background. The exon-6 specific genotyping primers, *Cul3*-ex6-Fwd: CTGCATGAAATGTGTCTTTAACTGT and *Cul3*-ex6-Rev: CCACCCCACTGCTAACTAGG, were used for PCR genotyping followed by Sanger sequencing. The Heterozygous *Cul3* founder mouse harboring 1bp-insertion in exon6 was expanded via breeding with wild-type (WT) C57BL/6N mice. The carrier male *Cul3^+/-^* mice were bred for at least four generations before starting any experiments to eliminate off-target effect of CRISPR. *Cul3* heterozygous (*Cul3^+/-^*) mice were viable, reached normal lifespan, and were fertile irrespective of sex. Animals were housed in groups of 4 animals per cage and kept on a 12 hour light/dark cycle (lights on at 7:00 am), with food and water available *ad libitum*. Embryonic time points were determined by plug checks, defining embryonic day (E) 0.5 as the morning after copulation. All experiments were approved by the Institutional Animal Care and Use Committees (IACUC) of the University of California San Diego (protocol number-S17029).

### Analysis of CRISPR off-target genes

The sgRNA was designed to target a patient-specific mutation in *Cul3*. In order to reduce off-target effects, we used Cas9 protein, which have much shorter half-life (about 24 hrs in the cells) than Cas9-plasmid or Cas9-mRNA. In addition, we back-crossed mutation carrier animals for four generations with C57BL6N animals to eliminate the remaining off targets. Two different software were used to predict off-target effects of SgRNA were used (https://chopchop.cbu.uib and https://crispr.cos.uni-heidelberg.de/). Top 10 off-target regions have mismatch of 3 base pairs or more, and they are present in exonic or intronic region of the genes, or and spliced ESTs sites (**Supplementary Table S12**). We also investigated the mRNA expression level of off-target genes by analyzing TPM values for the off-target genes in our transcriptomic data to make sure that the gene expression of the off-targets was not altered. We observed that only Cul3 was significantly downregulated in *Cul3^+/-^* mice, and all other genes had similar expression levels in both WT and *Cul3^+/-^* mice (**Supplementary Figure S14**).

### Mouse behavioral testing

*Cul3^+/-^* and WT littermates were used for behavioral experiments. Male and female mice were group housed post-weaning, segregated by sex, with unrestricted access to food and water, and were kept on a 12 h light/12 h dark cycle. The number of animals used in each behavior testing is mentioned in the figure legends. For all tests 1 hour of habituation to the room before each test was performed. *Cul3^+/-^* and their WT littermates were evaluated on the same day and by the same equipment, equipment was cleaned with 70% ethanol after each animal. Testing for open field, social preference, object recognition, repetitive behavior and social behavior were conducted during the light phase (10 am-5 pm) on different days. All testing equipment was positioned inside a photo light box that have dim overhead fluorescent lighting (14 lux). Tests were performed starting with the least aversive to most aversive task, and either scored automatically or by an experimenter blind to the genotype.

#### Developmental milestones

Development of reflexes and growth was evaluated as previously described ^32^. Reflexes were considered to be acquired only after they had been observed for 2 consecutive days.

#### Open Field

The apparatus measured approximately 43 x 43 x 33 cm (width x depth x height), the center was defined as the middle 26 x 26 cm. Mice were tested in a white open field for a total of 30 minutes per animal. The behavior and time spent in the center *vs* boundaries was recorded for 30 minutes by video camera. The videos were analyzed by MATLAB software and scored for 10, 20 and 30 minutes.

#### Novel object recognition

On day 1 of the experiment, mice were familiarized with the training room (1h) in their home cages. During the first day, the mice explored an empty chamber for 5 min (plastic chamber, 61×42×22 cm). On testing day, the animals were placed in the testing chamber for 1 min to re-habituate. Then two identical objects (plastic figures) were placed into the testing chamber. The mouse was placed in the chamber for 15 min to explore the objects. After a 3h delay, the mouse was placed in the testing chamber with one object identical to the previous object, and a new object with different shape, texture and color. Over a 15-min time period, the relative amount of time the animal spends exploring the new object compared to the familiar one was quantified by MATLAB software.

#### Repetitive behavior

Each mouse was placed in a novel test cage with no bedding or food. The animal spent a total of 20 minutes in novel test cage (10 minutes for habituation, 10 minutes for observation). The last 10 minutes of repetitive behavior was scored for the following categories: self-grooming, circling, isolation, or another unusual behavior. The scoring was done manually by an experimenter blind to the genotype.

#### Social preference test

The three-chamber test was done in a social box and grids were purchased from Harvard Apparatus (Catalog #76-0673, #76-0674). The test was performed as previously described ^33^. One day before testing mice were habituated to the three-chamber apparatus, and were allowed to explore the apparatus for 10 min, with empty grid at each side of the chamber. A plastic dish was placed on top of each grid to prevent climbing to the top. On test day, the test mouse was then placed in the center chamber and was free to explore the chambers for 10 min in each of the following phases: the first phase two identical non-social stimuli (Red lego blocks); the second phase non-social stimulus (yellow, gray and green Lego Blocks), and a social stimulus (age and sex matched mouse); the third phase familiar mouse and a novel mouse. The familiar mouse is the same mouse used in the second phase. The chamber was cleaned with 70% ethanol between phases. The chamber time and sniffing time was scored for the intervals of 10 min. Manual (blind analysis) and automated (MATLAB Software) methods of scoring chamber time and cylinder sniffing were used.

### Magnetic Resonance Imaging (MRI) of adult and early postnatal brains

The cohort used for behavioral testing was subsequently used for adult brain MRI study.

#### Perfusion

Mice undergone behavioral testing (*Cul3^+/-^*: 10 male, 10 female; WT: 8 male, 10 female) were anesthetized with a ketamine/xylazine mix (10mg/kg) and intracardially perfused with 30mL of 0.1 M PBS containing 10 U/mL heparin (Sigma) and 2mM ProHance (a Gadolinium contrast agent), followed by 30 mL of 4% paraformaldehyde (PFA) containing 2mM ProHance. After perfusion, mice were decapitated and skin, lower jaw, ears and cartilaginous nose tip were removed as previously described ^34, 35^. The brain within the skull was incubated in 4% PFA containing 2mM ProHance overnight at 4 degrees Celsius then transferred to 0.1M PBS containing 2mM ProHance and .02% sodium azide for at least 7 days prior to MRI scanning.

#### Imaging

After perfusion, a multichannel 7.0 Tesla MRI scanner (Agilent Inc., Palo Alto, CA) was used to image the brains within their skulls. Sixteen custom-built solenoid coils were used to image the brains in parallel. In order to detect volumetric changes via anatomical imaging, the following parameters were used: T2-weighted, 3-D fast spin-echo sequence, with a cylindrical acquisition of k-space, a TR of 350 ms and TEs of 12 ms per echo for 6 echoes, field-of-view equal to 20 × 20 × 25 mm^3^ and matrix size equal to 504 × 504 × 630 mm^3^. These parameters output an image with 0.040-mm isotropic voxels. The total imaging time was 14 h.

#### Analysis

All images were registered through iterative linear and nonlinear registrations to create a consensus average brain image. Deformation fields were computed which describe the differences between each individual and the average. The Jacobian determinants of the deformation fields were then calculated as measures of volume at each voxel ^36^. Structure volume was also calculated by warping a pre-existing classified MRI atlas onto the population atlas, which allowed for the calculation of 182 structures ^37–39^. Multiple comparisons were controlled for using the false discovery rate (FDR) ^40^.

### Young mice (P7) - 56um DTI

#### Perfusion

For younger mice, a similar perfusion protocol was followed with minor modifications. 15ml instead of 30ml of perfusion solution was used at both stages of the intracardial perfusion. Additionally, the skin and other skull structures were not removed following decapitation to prevent possible damage to the skull and brain of the neonatal mouse.

#### Imaging

For younger brains, diffusion tensor imaging was used to optimize gray/white matter contrast, as brains have yet to undergo large scale myelination in development. The diffusion sequence is a 3D diffusion-weighted FSE, with TR= 270 ms, echo train length = 6, first TE = 30 ms, TE = 10 ms for the remaining 5 echoes, one average, FOV = 25 mm × 14 mm × 14 mm, and a matrix size of 450 × 250 × 250, which yielded an image with 56 μm isotropic voxels. One b=0 s/mm 2 image is acquired and 6 high b-value (b = 2147 s/mm 2) images were acquired at the following directions (1,1,0), (1,0,1), (0,1,1), (-1,1,0), (-1,0,1) and (0,1,-1) corresponding to (G x, G y, G z). Total imaging time was ∼ 14 hours.

#### Analysis

First, the six high b-value images were averaged together to make a high contrast image necessary for accurate image registration. Then, all average high b-value images were linearly and nonlinearly registered together to create a consensus average brain. Deformation fields, which describe the deformations needed to take each individual mouse anatomy into the consensus average space, were calculated, and the Jacobian determinant of those deformation fields were then calculated as measures of volume at each voxel. Structure volume was also calculated by warping a pre-existing classified MRI atlas onto the population atlas, which allowed for the calculation of 56 structures. Multiple comparisons were controlled for using the false discovery rate (FDR) ^40^.

To calculate the change in region-specific volumes for a larger regions such as somatosensory cortex, cerebellum and entorhinal cortex **(Figure 2E)**, the sum of the volumes of all sub-regions comprising each of these larger regions was calculated for each animal. Then, the mean, standard deviation, effect size and % difference were calculated for WT and *Cul3*+/- groups. The two tailed T-test was used for comparing the statistical differences.

### Immunohistochemistry

Mice were deeply anesthetized by xylazine and ketamine and were perfused with 4% paraformaldehyde (PFA). Whole brain tissues were extracted and dehydrated in a 30% sucrose solution at 4°C overnight. The 30 μm thick brain slices were prepared using a cryostat (Leica, Germany). Slices were rinsed in 0.01 M PBS, permeabilized with 0.1%TritonX-100 in PBS and blocked in 3% FBS (with 0.1% Triton X-100) for 1 h. Primary antibody against NeuN (mouse 1:1000, Millipore, USA), BRN2 (rabbit 1:1000, cell signaling technology, USA) was added for overnight incubation at 4°C. Donkey anti-rabbit Alexa 555 and anti-mouse Alexa 488 secondary antibody (Life Technology, Camarillo, CA, USA) was added at room temperature for 2 h incubation. After PBS rinsing, DAPI was added for nuclear staining. Slices were mounted using ProLong Gold antifade mountant (Invitrogen). Fluorescent images were taken on Leica SP8 confocal microscope using 10x inverted objective and quantification was done using ImageJ (NIH). To examine cortical layering, the length of the layer containing NeuN-positive cells within the somatosensory cortex was measured (N=6 littermate animals per genotype). For the quantification of mature neurons in layer II-IV, the number of total NeuN-positive cells and BRN2-positive cells in II-IV cell layers was performed within the somatosensory cortex and normalized to the area used (N=6 littermate animals per genotype).

### Primary cortical neuron culture

Cortices were dissected from WT and *Cul3^+/^*^-^ mouse embryonic brains at E17.5. Cortex from each brain was cultured individually. Dissociation was initiated by incubating the dissected cortices in 1ml of Accumax (Innovative Cell Technologies Inc) for 30 min at 37°C followed by 5 min incubation in 10 mg/ml DNaseI (Sigma Aldrich), the dissociated cells were gently triturated with fire-polished glass Pasteur pipette to make single cell suspension, and then passed through 40-micron nylon filter to remove any non-dissociated tissue. Cells were counted with cell counter (BioRad) and 1×10^5^ cells were seeded onto glass coverslip coated with 0.01 % P-L-ornithine and 5ug/mL mouse natural laminin in 24-well plates. The plating media contained Neurobasal medium, 2% B27 supplement, 10% horse serum, 1% penicillin/streptomycin, and 2 mM L-glutamine (Invitrogen). After 12 hr, media was changed to serum-free feeding media containing Neurobasal medium, 2% B27 supplement, 1% penicillin/streptomycin, and 2 mM L-glutamine. At DIV2-4, cultures were treated with 1mM Cytosine b-D-arabinofuranoside hydrochloride (Ara-C) (Sigma Aldrich) to inhibit glial cell proliferation. The cells media were half replaced with new media containing AraC every four days. Cultures were maintained at 37°C and 5% CO_2_. All media components were from GIBCO unless otherwise specified.

### Immunocytochemistry

Coverslips containing primary cortical neurons were fixed in 4% paraformaldehyde (PFA) for 15 min and washed three times with PBS. Permeabilization and blocking was performed with 3% Bovine Serum Albumin (BSA, Sigma-Aldrich), 0.1% Triton X-100 (Sigma Aldrich) in PBS for one hour at room temperature. The coverslips were then incubated overnight at 4°C with primary antibodies diluted in solution containing 3% BSA. PBS was used to wash the primary antibodies and the coverslips were incubated with secondary antibodies in solution containing 3% BSA for 1.5 h at room temperature. The following primary antibodies were used for immunostaining: MAP2 (rabbit 1:1000, Cell Signaling Technology); Rhodamine Phalloidin (Cytoskeleton). Alexa Fluor Dyes (Abcam) were used at 1:1000 dilution as secondary antibodies. Nuclei were visualized with DAPI (1:25000, Life Technologies). Slides were mounted using ProLong Gold antifade reagent (Invitrogen) and analyzed under a fluorescence microscope (Leica SP8). Image analysis was performed with ImageJ software

### Morphological Analysis

Images were taken with Leica SP8 confocal microscope with oil-inverted 40x objective and processed and analyzed with ImageJ 1.5 software. For soma area calculation, the perimeter of the Map2-positive cell bodies were manually outlined and measured. For total dendrite length analysis, the simple neurite tracer plugin for ImageJ 1.5 was used. Map2-positive neurites from each neuron were traced and the dendrite length was calculated by adding individual lengths for every neuron. F-actin positive puncta on dendrites was calculated by visually counting all protrusions from a dendrite within a 15–25 mm distance starting at a secondary branch point. One to three dendritic segments were analyzed per neuron. The number of F-actin puncta was normalized by dendritic length, similar region were also used for quantification of F-actin intensity using Image J 1.5. Image analysis and quantification was performed by the trained experimenter blind to the genotype.

### Multi-electrode array (MEA) recording

Primary cortical neurons from WT and *Cul3^+/-^* E17.5 embryos were cultured on 12-well Axion Maestro multielectrode array (MEA) plates (Axion Biosystems, Atlanta, GA, USA). Each 12-well MEA plate from Axion Biosystems was coated with 100 μg/mL poly-L-ornithine and 10 μg/mL laminin for 24 h before seeding. 1.5×10^6^ neurons were plated on the coated MEA plates and a half-medium change was performed every other day using neurobasal+B27 feeding medium. Spontaneous spike activity was recorded on 8 DIV. Recordings were performed at 37°C using a Maestro MEA system and AxIS Software Spontaneous Neural Configuration (Axion Biosystems). Briefly, Spikes were detected with AxIS software using an adaptive threshold and then analyzed using Axion Biosystems’ Neural Metrics Tool. Spike time stamps were exported to Neuroexplorer (Axion Biosystems) for the creation of raster plots. Bright-field images were captured to assess cell density and electrode coverage.

### Pharmacological treatment of cortical neurons with Rhosin

For morphological phenotype rescue experiments, cultured cortical neurons were grown in Rhosin-treated neurobasal media. Rhosin (Tocris) was added to the final concentration of 10µm to the media during at 2 DIV stage. The same amount of Rhosin was added every 4 days during all subsequent media changes. The cells were grown for 14 days, at which dendrite staining was carried out with Map2 antibody as mentioned in immunocytochemistry section. An equivalent amount of vehicle (Dimethylsulfoxide, DMSO) was added to a final concentration of 0.001% to growth media to obtain vehicle-treated WT and Cul3^+/-^ cortical neurons. For MEA phenotype rescue experiments, the media changes were performed every 3^rd^ day. The cells were grown for 8 days, at which MEA recording and image acquisition was carried out. After completion of the experiment the cells were extracted and lysed to quantify total protein within each well.

### TUNEL Assay

We used a Terminal Deoxynucleotidyl Transferase (TdT) dUTP Nick-end Labeling (TUNEL) assay to measure the apoptosis of the primary cortical neurons. The assay was performed using APO-BrdU TUNEL assay kit (Invitorgen) as described by manufacturer. In brief, primary cortical neurons were fixed with 4%PFA followed by O/N incubation in 70% ethanol at -20°C followed by PBS washing. A freshly prepared TUNEL reaction buffer (50 μL per sample) was added at 37°C incubation for 1 h. After rinsing with the rinse buffer, the Propidium Iodide (PI) was used for nuclei staining. The cells were acquired in BD accuri C6 (BD, Franklin Lakes, NJ) flow cytometer. All the acquired cells were considered as population of interests for further analysis. Respective single color-stained samples were used to correct color compensation overlapping between two channels. The percentage of TUNEL-positive and PI-positive cells were analyzed with FlowJo software (FlowJO LLC, Ashland, OR) with respect to the corresponding unstained samples. The percentage change in dual positive cells, *i.e.* PI- and BrdU-488 positive cells was calculated.

### RNA isolation for bulk RNA sequencing and qPCR

Total RNA was extracted at three developmental time periods (embryonic E17.5, early postnatal (day 7) and adult (4-6weeks)) from three brain regions (cerebral cortex, hippocampus, and cerebellum) of WT and *Cul3^+/-^* mice using the QIAGEN RNAeasy isolation kit (QIAGEN) following manufacturer’s instructions. RNA sequencing was performed using equal input amount of total RNA for each sample. RNA samples were ribodepleted using Ribo-Zero rRNA Removal Kit (Illumina) and library preparation was performed using the True-Seq Stranded Total RNA kit for Illumina Sequencing according to the manufacturer’s instructions. Paired-end RNA sequencing (2×150bp) was performed on an Illumina HiSeq4000 to an average depth of 40M reads per sample.

For Quantitative RT-PCR (qPCR) experiments (**Figure 1e**), cDNA was synthesized starting from 100ng of total RNA with the SuperScript III First-Strand Synthesis kit and oligo dT (Invitrogen). The qPCR was performed using the CFX96 Touch™ Real-Time PCR Detection System (Bio Rad) using Power SYBR Green PCR Master Mix (Applied Biosystems). The following primers were used for Cul3 - Cul3_Fwd: TCAAACAGTTGCAGCCAAAC and Cul3_Rev: GAATCGAGCCTTCAGTTGCT. GAPDH was used as housekeeping gene for the normalization, the following primers were used for GAPDH - GAPDH_Fwd: AGGTCGGTGTGAACGGATTTG and GAPDH_Rev: TGTAGACCATGTAGTTGAGGTCA. Fold change in expression was calculated using the ΔΔ^Ct^ method. Data is presented as levels of Cul3 normalized to GAPDH levels.

### RNA-seq Data Processing Pipeline

All 108 FASTQ files (36 embryonic, 36 early postnatal and 36 adult) (**Supplementary Figure S6**) were run through a unified paired end RNA-Seq processing pipeline. Pipeline source code can be found on https://github.com/IakouchevaLab/CUL3. All fastqs were trimmed for adapter sequence and low base call quality (Phred score < 30 at ends) using Cutadapt (v1.14). Trimmed reads were then aligned to the GRCm38.p5 (mm10) reference genome via STAR (2.5.3a) using comprehensive gene annotations from mouse Gencode (v16). Gene-level quantifications were calculated using RSEM (v1.3). Quality control metrics were calculated using RNA-SeQC (v1.1.8), featureCounts (v1.6.), PicardTools (v2.12), and Samtools (v1.3) (**Supplementary Table S13, Supplementary Figure S7**).

### RNA-Seq Quality Control, Normalization and Differential Gene Expression Analysis

RNA-Seq Quality Control and Normalization Expected counts were compiled from gene-level RSEM quantifications and imported into R for downstream analyses. Expressed genes were defined as genes with TPM > 0.1 in at least 80% of samples from each genotype (WT and *Cul3^+/^*). A total of 17,363; 18,656; and 18,341 expressed genes from embryonic cortex, cerebellum, and hippocampus, respectively; 18,921; 18,276; and 18,396 expressed genes from early postnatal cortex, cerebellum, and hippocampus, respectively; and 18,061; 17,835; and 17,654 expressed genes from adult cortex, cerebellum, and hippocampus, respectively, were used in the downstream analysis using the mouse Gencode (v16) annotation gtf file. Only expressed genes were included into the analysis.

To account for possible batch effects, batch effect removal was performed using EDASeq R package ^41^ and “remove unwanted variation” (RUVs) package in RUVseq ^42^. Initial normalization of count data was done using upper quartile normalization ^41^, followed by RUVseq. The parameter k in RUVseq was set to the minimum value that separated the samples based on genotypes on the PCA plot (maximum k=10). RUVs with different k values ranging from 3 to 8 were used after defining groups based on genotype (i.e. WT vs *Cul3^+/-^*). Differential gene expression analysis was performed with edgeR ^43^ using the negative binomial GLM approach and by considering design matrix that includes both covariates of interest (Genotype and Gender) and the factors of unwanted variation. Genes with a false discovery rate (FDR ≤ 0.1, Benjamini-Hochberg multiple testing correction) were considered significant and used for further downstream processing and analysis.

Differentially expressed gene datasets for all regions (CX, CB and HP) across individual developmental periods, and all periods (E17.5, P7 and P35) across individual brain regions, were considered for period- and region-wise P-value combination analysis. We performed a total six P-value combination analysis that included both, period- and region-wise comparisons. Preprocessing of the DEG datasets was done by selecting differentially expressed genes with similar trend (i.e. either up- or downregulated in mutant *vs* WT) based on log fold change expression, for respective regions-wise or period-wise comparisons. The Fisher P-value combination method with metaRNASeq R package ^44^ was used to combine the P-values for these sets of genes, following similar trend in the log fold change expression for each comparison. The combined FDR-corrected P-value for each set of genes was obtained using Benjamini-Hochberg method. Genes with combined FDR-corrected P-value of ≤ 0.1 in each region- and period-wise comparison were hierarchically clustered. We obtained a total 21 clusters from period- and region-wise analyses combined. GO terms enrichment was performed using gProfiler ^45^ for all gene clusters, resulting in around 1700 GO terms. Log fold GO enrichment was performed for all GO terms, as well as for shared by two or more clusters GO terms. These shared GO terms were subsequently hierarchically clustered to produce log fold GO term enrichment Figure (**Figure 4d**).

### Gene Ontology enrichment analysis

Enrichment of Gene Ontology terms (GO) biological process (BP) and molecular function (MF) was performed using gProfiler R package ^45^. Background was restricted to the expressed set of genes by period (embryonic, early postnatal, adult) or region (cortex, cerebellum and hippocampus) whenever appropriate. Ordered query was used, ranking genes by FDR-corrected P-value.

### Enrichment analysis of ASD-relevant gene sets

Enrichment analyses were performed using several established, hypothesis-driven gene sets including syndromic and highly ranked (1 and 2) genes from SFARI Gene database (https://gene.sfari.org/database/gene-scoring/); pre- and post-synaptic genes from SynaptomeDB ^46^; genes with loss-of-function intolerance (pLI) > 0.99 as reported by the Exome Aggregation Consortium ^47^; highly constrained genes ^48^; and FMRP targets ^49^. Fisher exact test was used to calculate the enrichment of significantly differentially expressed genes for each curated gene set. The background lists were the union of all analyzed genes. Significance values of the results were corrected for multiple hypothesis testing using Benjamini-Hochberg method.

### Quantitative TMT-Mass Spectrometry

#### Sample preparation

TMT mass-spectrometry experiments were performed on the WT and *Cul3^+/-^* cerebral cortex and cerebellum at three developmental periods (embryonic, early postnatal and adult). Tissues were collected from the remaining half of hemisphere of the brains used for RNA-seq experiments (except for embryonic and adult cerebellum), and snap-frozen in liquid nitrogen. After collection of the entire set, the tissues were lysed in RIPA buffer (20 mM Tris, pH 7.4, 150 mM NaCl, 1 mM EDTA, 1% sodium deoxycholate and 1% Triton X-100) supplemented with 1xEDTA-free complete protease inhibitor mixture (Roche) and phosphatase inhibitor cocktails-I, II (Sigma Aldrich). The lysates were centrifuged at 16,000xg at 4°C for 30min, and the supernatants were collected. Protein concentration was quantified by modified Lowry assay (DC protein assay; Bio-Rad). Lysates were subjected to methanol-chloroform precipitation, resuspended in 8M urea in 50 mM TEAB, reduced (10 mM TCEP at room temperature for 25 min) and alkylated (50 mM chloroacetamide at room temperature in the dark for 20 min). After another round of methanol-chloroform precipitation, pellets were dissolved by adding 6M urea in 50 mM TEAB, and the protein concentration was estimated by BCA assay (Thermo, 23225). LysC/Tryp (Promega, V5073) was added by 1:25 (w/w) ratio to the proteins. After 3-4 h of incubation with 850 rpm shaking at 37°C, reaction mixture was diluted with 50 mM TEAB for urea to be less than 1 M. After the o/n digestion, peptide concentration was estimated by colorimetric peptide BCA assay (Thermo, 23275), and the peptides were labelled with TMT 10-plex reagents (Thermo, 90110) for one hour, followed by 15 min quenching with hydroxylamine according to the manufacturer’s protocol. Equal amounts of reaction mixtures for each channel were pooled together and dried using SpeedVac. Dried peptides were resuspended in 0.1% TFA and fractionated using Pierce™ High pH reversed-phase peptide fractionation kit (Thermo, 84868) and then dried in SpeedVac.

#### Mass spectrometry

Each fraction was dissolved in buffer A (5% acetonitrile, 0.1% formic acid) and injected directly onto a 25 cm, 100μm-ID column packed with BEH 1.7μm C18 resin (Waters). Samples were separated at a flow rate of 300 nl/min on nLC1000 (Thermo). A gradient of 1–30% B (80% acetonitrile, 0.1% formic acid) over 160 min, an increase to 90% B over another 60 min and held at 90% B for a final 20min of washing was used for 240 min total run time. Column was re-equilibrated with 10μL of buffer A prior to the injection of sample. Peptides were eluted directly from the tip of the column and nanosprayed directly into the mass spectrometer Orbitrap Fusion by application of 2.8 kV voltage at the back of the column. Fusion was operated in a data dependent mode. Full MS1 scans were collected in the Orbitrap at 120K resolution. The cycle time was set to 3s, and within this 3s the most abundant ions per scan were selected for CID MS/MS in the ion trap. MS3 analysis with multi-notch isolation (SPS3) ^50^ was utilized for detection of TMT reporter ions at 60K resolution. Monoisotopic precursor selection was enabled, and dynamic exclusion was used with exclusion duration of 10s.

#### Protein identification and quantification

Tandem mass spectra were extracted from the .raw files using Raw Converter ^51^ with monoisotopic peak selection. The spectral files from all fractions were uploaded into one experiment on Integrated Proteomics Applications (IP2, Ver.6.0.5) pipeline. Proteins and peptides were searched using ProLuCID ^52^ and DTASelect 2.0 ^53^ on IP2 against the UniProt reviewed Mus musculus protein database with reversed decoy sequences (UniProt_mouse_reviewed_contaminant_05-25-2018_reversed.fasta from IP2 standard database). Precursor mass tolerance was set to 50.0 ppm, and the search space allowed both full- and half-tryptic peptide candidates without limit to internal missed cleavage and with a fixed modification of 57.02146 on cysteine and 229.1629 on N-terminus and lysine. Peptide candidates were filtered using DTASelect parameters of -p 2 (proteins with at least two peptides are identified) -y 1 (partial tryptic end is allowed) --pfp 0.01 (protein FDR < 1%) -DM 5 (highest precursor mass error 5 ppm) -U (unique peptide only). Quantification was performed by Census ^54^ on IP2.

### Differential protein expression analysis

Proteomics data was first summarized to peptide level by adding up the intensities of constituting spectra. Quantitation results from different TMT runs were merged and normalized using the pooled samples channel which was present in all runs. For each peptide, multiple measurements from the same mouse were collapsed to one measurement by taking the sum of all measurements. Batch effects from the summarized protein-level data for each dataset were removed using ComBat ^55^. The data was then log_2_ transformed. Differentially expressed proteins were identified using function lmFit in limma ^56^ followed by eBayes moderation of standard errors ^57^. Resulting P-values were FDR-corrected using the Benjamini-Hochberg method to control for multiple comparisons.

### Weighted protein co-expression network analysis

We used Weighted Protein Co-expression Network Analysis (WPCNA) ^58^ to define modules of co-expressed proteins from proteomics data. Proteomics data was first summarized to protein level by adding up the channel intensities of constituting peptides for each of six datasets derived from two brain regions (cortex and cerebellum) and three developmental periods (embryonic, early postnatal and adult) (**Supplementary Figure S9**). Batch effects from the summarized protein-level data for each dataset were removed using ComBat ^55^, followed by log_2_ transformation. Modules were estimated using the blockwiseModules function with the following parameters: corType=bicorr; networkType=signed; pamRespectsDendro=F; mergeCutHeight=0.1. Some parameters were specific to each dataset. For embryonic cortex: power=18; deepSplit=0; minModuleSize=100; for early postnatal cortex: power=18; deepSplit=0; minModuleSize=100; for adult cortex: power=18; deepSplit=0; minModuleSize=20; for embryonic cerebellum: power=22; deepSplit=0; minModuleSize=40; for early postnatal cerebellum: power=26; deepSplit=0; minModuleSize=150; and for adult cortex: power=12; deepSplit=2; minModuleSize=40. Module eigengene-genotype associations were calculated using linear regression model. Significance P-values were FDR-corrected to account for multiple comparisons. Genes within each module were prioritized based on their module membership (kME), defined as correlation to the module eigengene.

### Cul3 expression in human and mouse published datasets

Human RNA-seq data (as RPKM) for neocortex was downloaded from the BrainSpan Atlas of the Developing Human Brain (https://hbatlas.org/hbtd/basicSearch.pl). For the developmental trajectory, Neocortex (NCX) was selected and the expression of Cul3 was plotted for time points up to 37 years of age. The data was plotted as mean across samples, error bar represents standard deviation. For expression profile of Cul3 in mouse, the data from C57Bl6 mice were extracted from Gompers et al. ^59^ and Cul3 expression was plotted starting from E12.5 up to adult.

### Cell type enrichment analysis using mouse scRNA-seq data

Cell-type enrichment analysis for each protein co-expression module was performed using the Expression Weighted Cell Type Enrichment (EWCE) package in R ^60^. Cell type-specific gene expression data was obtained from single cell sequencing (scRNA-seq) studies of Postnatal (P0) mouse neocortex regions ^16^. The specificity metric of each gene for each cell type was computed as described ^60^. “Intermediate progenitors” cell type includes a union of SVZ 2 (migratory neurons), “Interneurons” cell type includes union of Interneuron 5P, 6P, 11P and 14P, “Layer V-VI” cell type includes 18P and 12P (excitatory neurons), and “Layer II-IV” cell type includes 15P and 1P (excitatory neurons). Enrichment was evaluated using bootstrapping. Z-score was estimated by the distance of the mean expression of the target gene set from the mean expression of bootstrapping replicates. P-values were corrected for multiple comparisons using FDR.

### Western Blotting

The small fraction of tissue lysates prepared for mass-spectrometry experiments were used for Western Blotting. On average 10-15ug of total protein from WT and *Cul3^+/-^* cerebral cortex, cerebellum or hippocampus were resolved by SDS-PAGE and transferred onto PVDF Immobilon-P membranes (Millipore). After blocking with 5% nonfat dry milk in 1X TBS with 0.1% Tween-20 (TBST) for 1h at room temperature, membranes were first probed overnight with the appropriate primary antibodies in 3% BSA in TBST, and then after 1h of incubation with corresponding host specific HRP-conjugated secondary antibody (Abcam). Membranes were developed using the EZ-ECL chemiluminescence detection kit (Denville Scientific). The following primary antibodies were used: anti-Cul3 (1:1000; Cell Signaling), anti-RhoA (1:1000; Cell Signaling), and anti-Gapdh (1:5000; Sigma Aldrich) as a loading control. Quantification was performed by densitometry with ImageJ software.

### Quantification and statistical analysis

The statistical analyses for the above experiments were performed using Prism software (GraphPad). Statistical tests, sample sizes and exact P-values are described in Figure legends. Significance was defined as p < 0.05(*), p < 0.01(**), or p < 0.001(***). Blinded measurements were performed for any comparison between control and *Cul3*^+/-^ genotypes.

### Data availability

Source RNA-seq data is available at GEO repository accession number GSE144046 (https://www.ncbi.nlm.nih.gov/geo/query/acc.cgi?acc=GSE144046).

Source proteomics data is available from the public repository MassIVE (Mass Spectrometry Interactive Virtual Environment), a part of the ProteomeXchange consortium, with the identifier MSV000084830 (and PXD017256 for ProteomeXchange) through the following link (https://massive.ucsd.edu/ProteoSAFe/dataset.jsp?accession=MSV000084830).

### Code availability

The code used for the RNAseq and proteomics analysis generated during this study is available from GitHub (https://github.com/IakouchevaLab/CUL3).

## Supplementary Figures titles and legends

**Figure S1.**
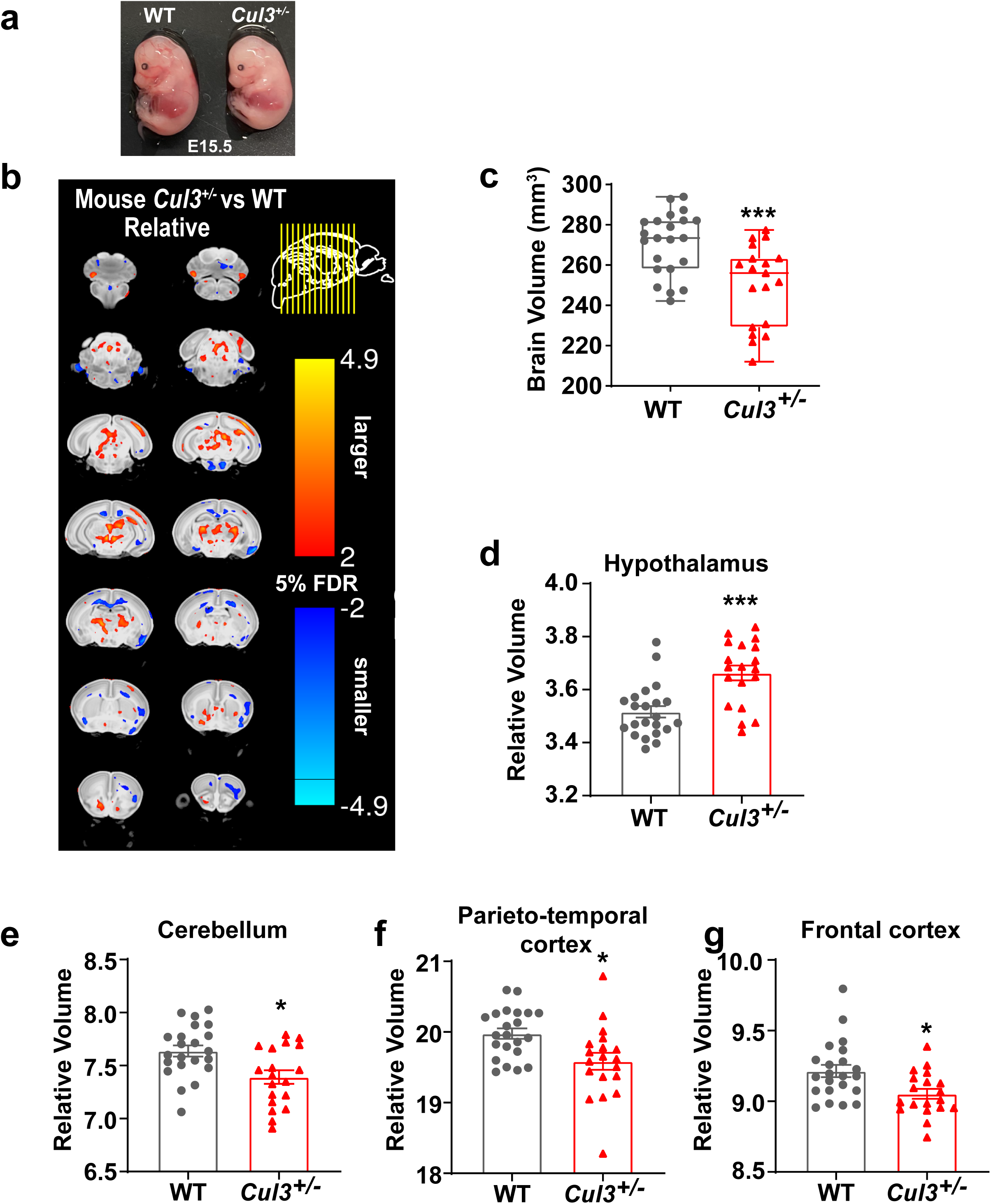
Altered brain morphology of *Cul3^+/-^* postnatal day 7 (P7) mice. **(a)** The representative images of WT and *Cul3^+/-^* E15.5 embryos, *Cul3^+/-^* embryos are smaller in size. **(b)** Voxel-wise analysis highlighting significant differences in relative volumes throughout the brain between WT and *Cul3^+/-^* mice with 5% false discovery rate (FDR). Scale bar 2-4.9 indicates decreasing FDR, where the value of 2 corresponds to 5% FDR, and positive or negative values are *vs* WT brain. **(c)** *Cul3^+/-^* mice have reduced absolute brain volume (***p<0.001) compared with WT mice (n=22 WT, 19 *Cul3^+/-^*). **(d)** MRI revealed significant increase in hypothalamus volume (\*\*\**p*<0.001). (**e-g**) MRI revealed significant reduction in relative volume (normalized by % total brain volume) of cerebellum (*p<0.05), parieto-temporal cortex (*p<0.05), and frontal cortex (*p<0.05); Dots represent individual samples; Two tailed T-test used for **c-g**; error bars represent mean ± SD.

**Figure S2.**
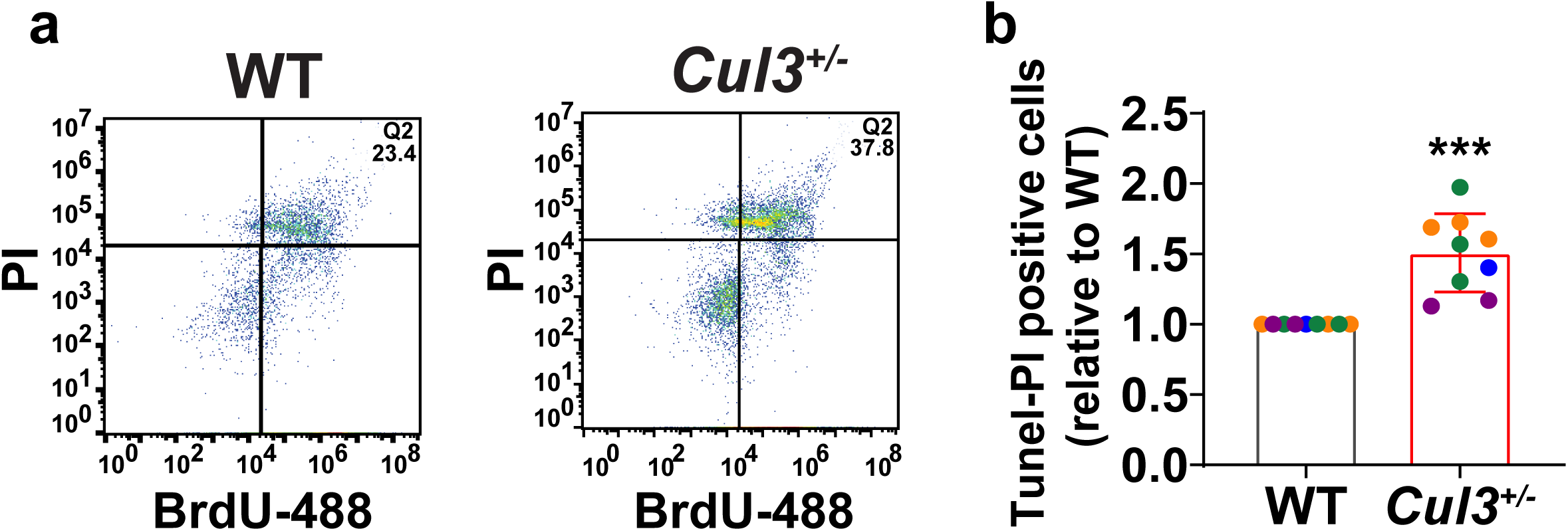
Increased apoptosis in *Cul3^+/-^* mice. **(a-b)** TUNEL assay on 14DIV primary cortical neurons, the dual positive neurons for PI and BrdU-488 were counted by flow cytometry; (Q4) were considered as apoptotic cells containing BrdU-488/PI+ cells (***p<0.001; n=9 for each genotype). Dots represent independent samples; two tailed T-test used for **b**; error bars represent mean ± SD

**Figure S3.**
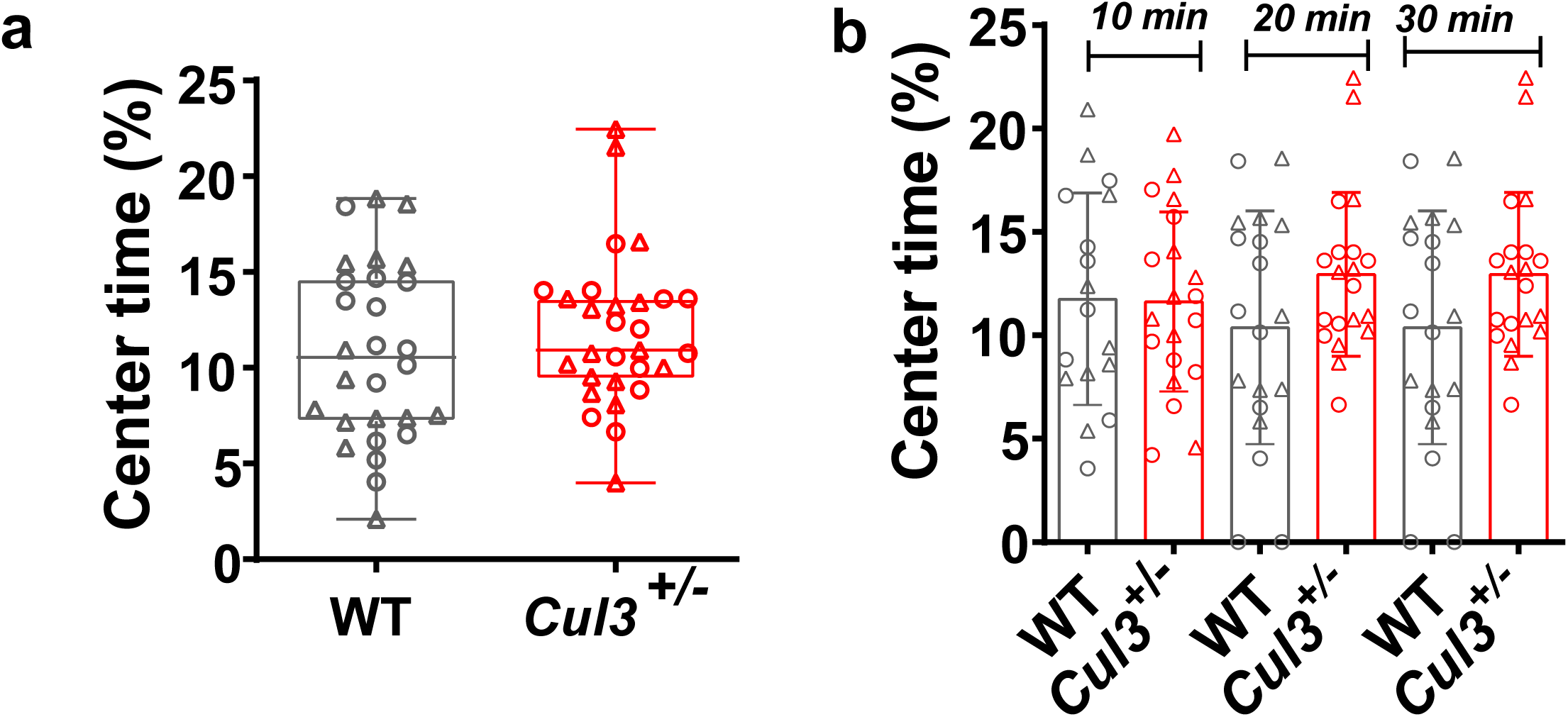
*Cul3^+/-^* mice do not display anxiety. **(a)** *Cul3^+/-^* mice display no preference for the center *vs* periphery in open field test. **(b)** 10 min time-bins showing no difference in % time spent in the center in any 10 min intervals. Dots represent individual animals; Two tailed T-test used for **a-b**; error bars represent mean ± SD. ○-female and Δ-male.

**Figure S4.**
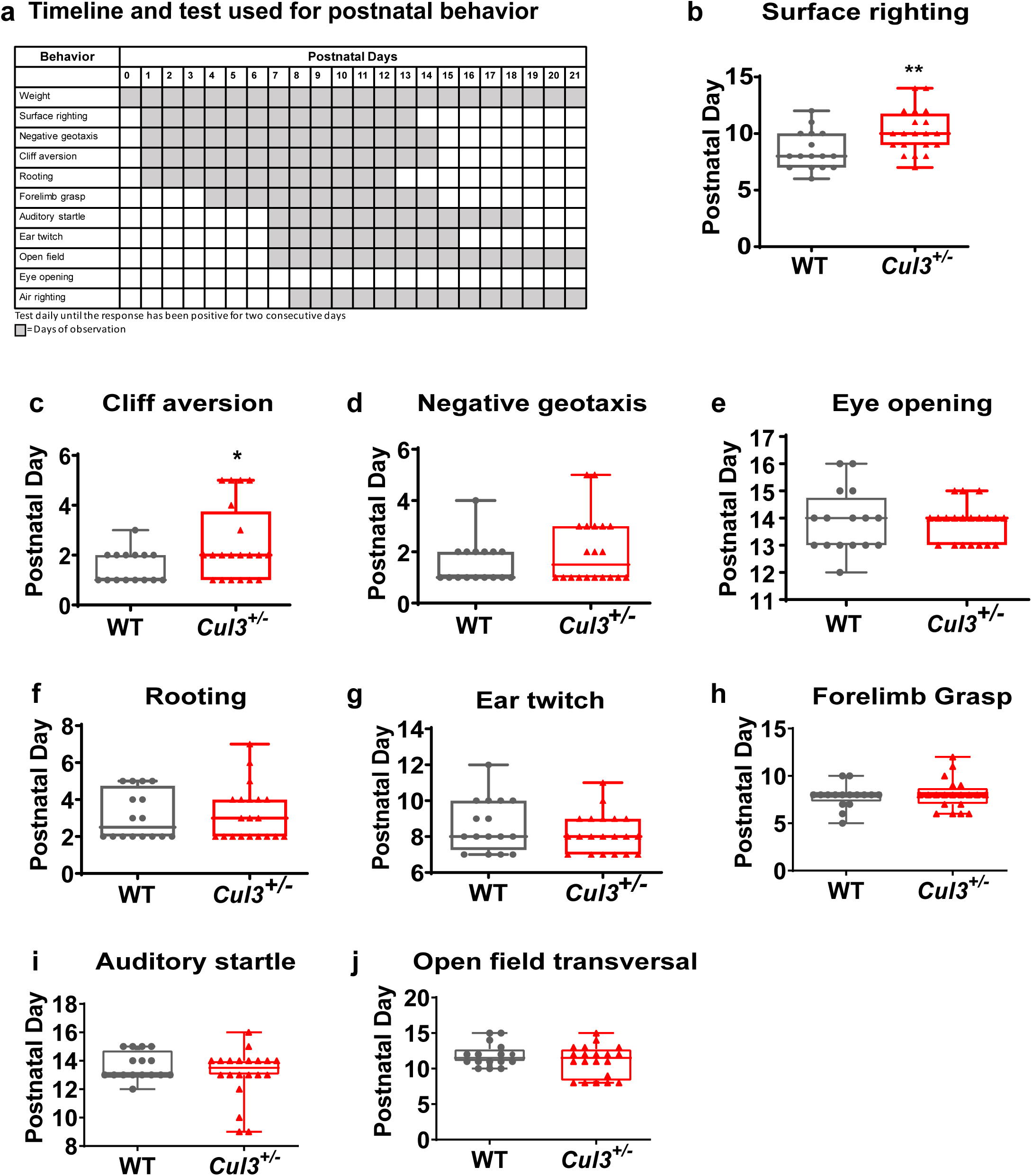
Analysis of developmental milestones and early postnatal behaviors in *Cul3^+/-^* mice. **(a)** The timeline used for the early postnatal behavioral assays. **(b)** The righting reflex is motor ability for a mouse pup to be able to flip onto its feet from a supine position, *Cul3^+/-^* mice display delayed righting reflex as compared to WT mice (n=17 WT, n=19 *Cul3^+/-^,* p<0.01). **(c)** *Cul3^+/-^* mice display significant vestibular imbalance. The pup’s eyes are still closed so fear is not the driving factor to turn away from the cliff’s edge (n=15 WT, n=20 *Cul3^+/-^*\*, p<0.05). **(d-k)** Both *Cul3^+/-^* and WT mice have no difference in other postnatal behaviors tasks. Dots represent independent animals; two tailed T-test used for **b-k**; error bars represent mean ± SD.

**Figure S5.**
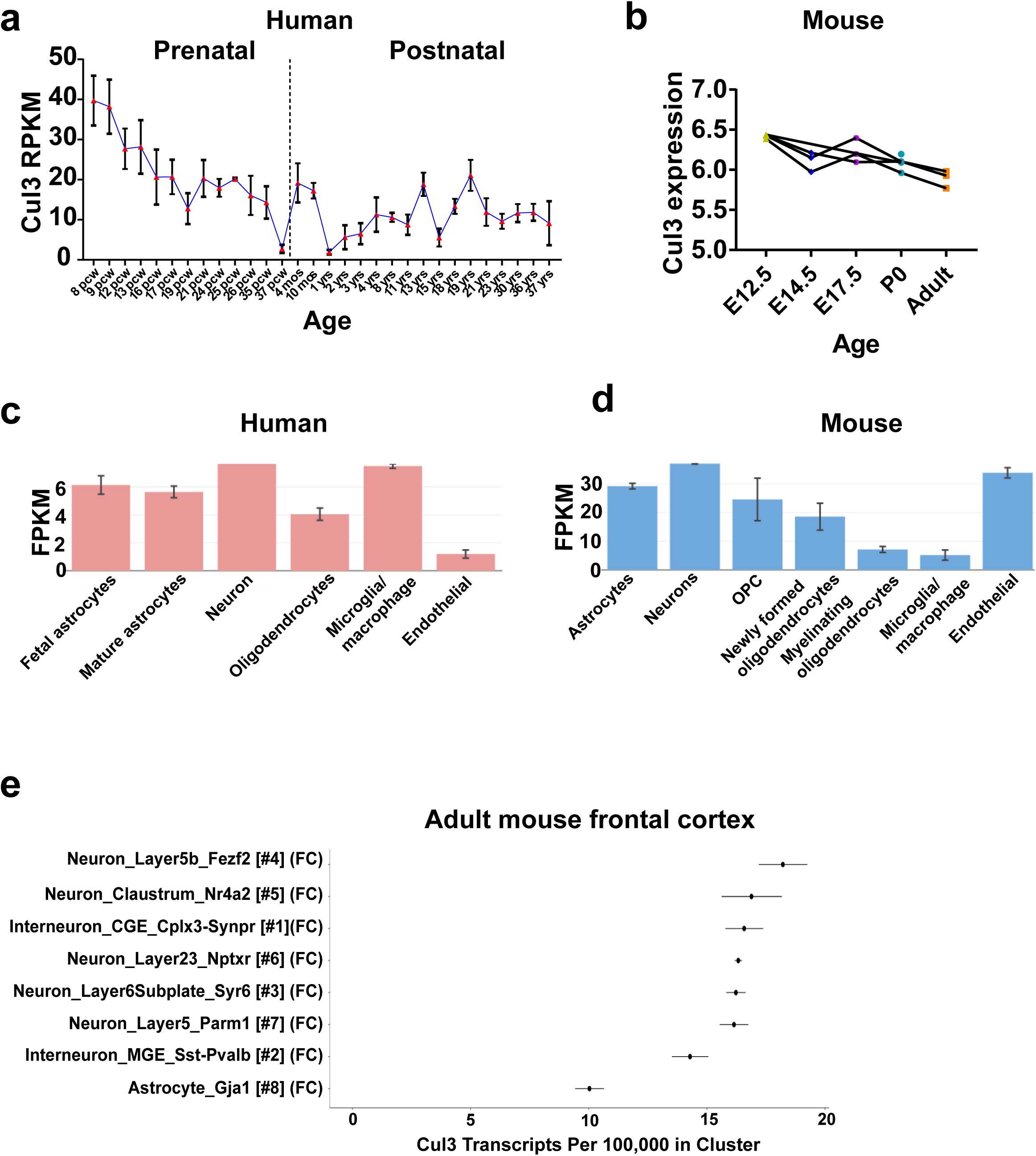
Cul3 expression in human and mouse across development using publicly available data. **(a)** Cul3 expression in the developing human neocortex extracted from Brainspan (https://www.brainspan.org/). **(b)** Cul3 expression in developing mouse brain based on Gompers datasets ^59^. (**c**) Cell-type specific expression of Cul3 in human brain (https://www.brainrnaseq.org/) ^61^. (**d**). Cell-type specific expression of Cul3 in mouse brain (https://www.brainrnaseq.org/) ^62^. **(e)** Cell-type specific expression of Cul3 in mouse brain based on DropViz (http://dropviz.org) ^63^.

**Figure S6.**
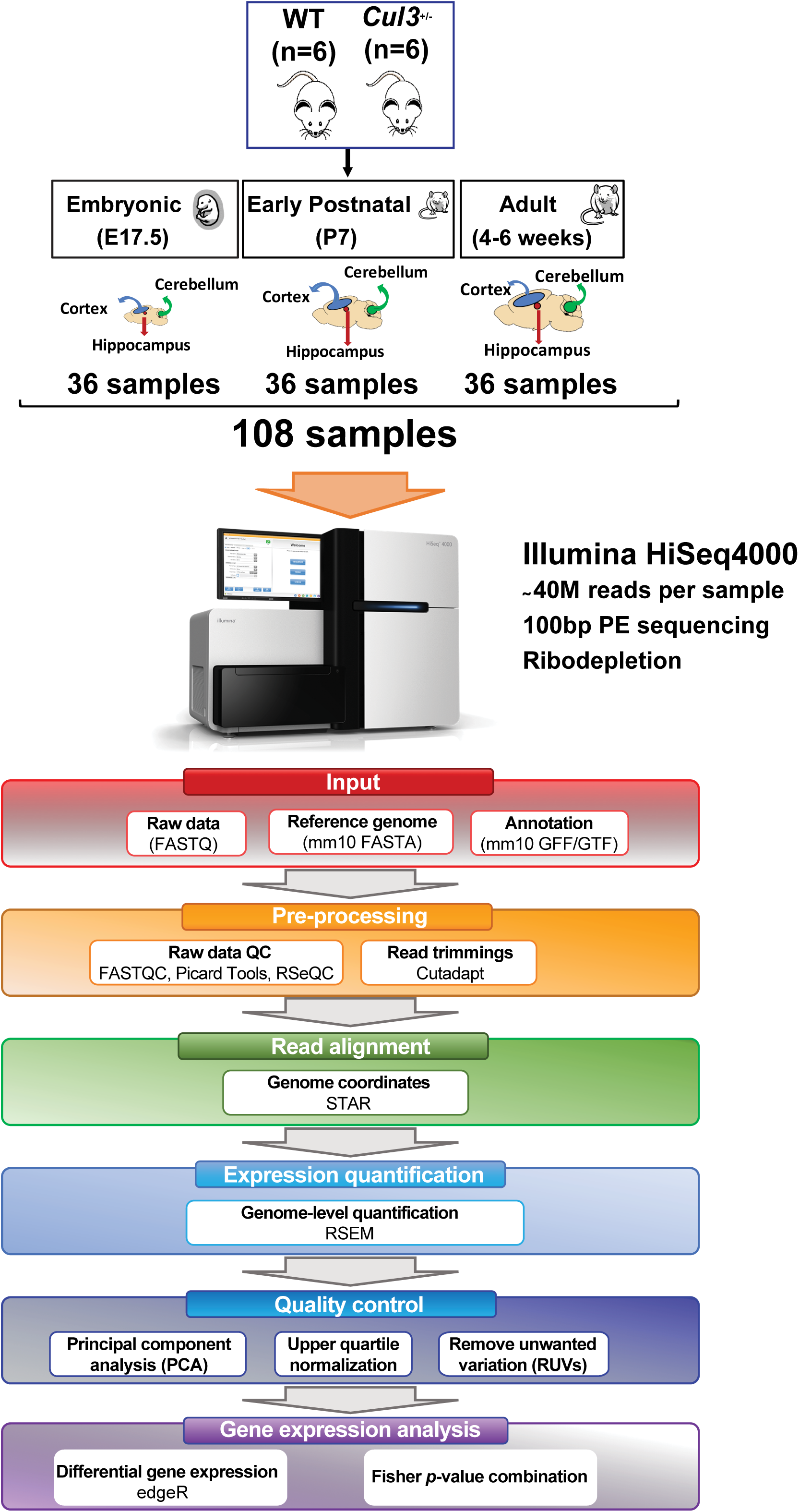
RNA-seq experimental design and data analysis workflow. A total of 108 transcriptomes have been bulk RNA sequenced in this study. A rigorous quality control including principal component analyses, upper quartile normalization, and removal of unwanted variables (RUV) was used for batch correction.

**Figure S7.**
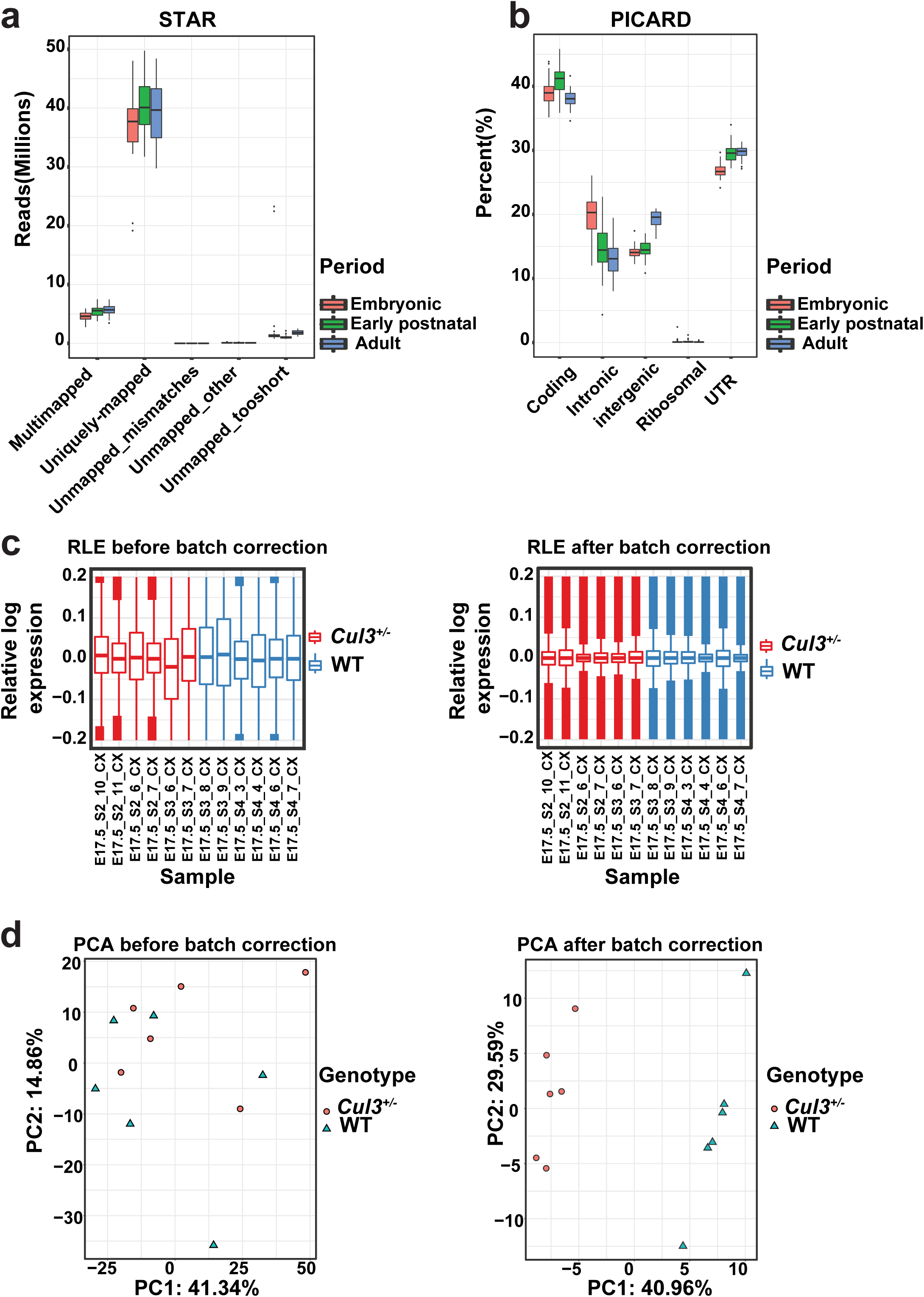
Quality control metrics for RNA-seq data. **(a)** Sequencing metrics from STAR (2.5.3a) for samples from three periods (embryonic E17.5, early postnatal P7 and adult). **(b)** Sequencing metrics from PICARD (v2.12) from three periods. **(c)** Relative log expression values are shown before (left panel) and after (right panel) batch correction using RUVseq for the embryonic cortex samples as an example. **(d)** First two principal components (PCs) of gene expression values calculated using “prcomp” function in R, are shown before (left panel) and after (right panel) batch correction using RUVseq for the embryonic cortex samples as an example.

**Figure S8.**
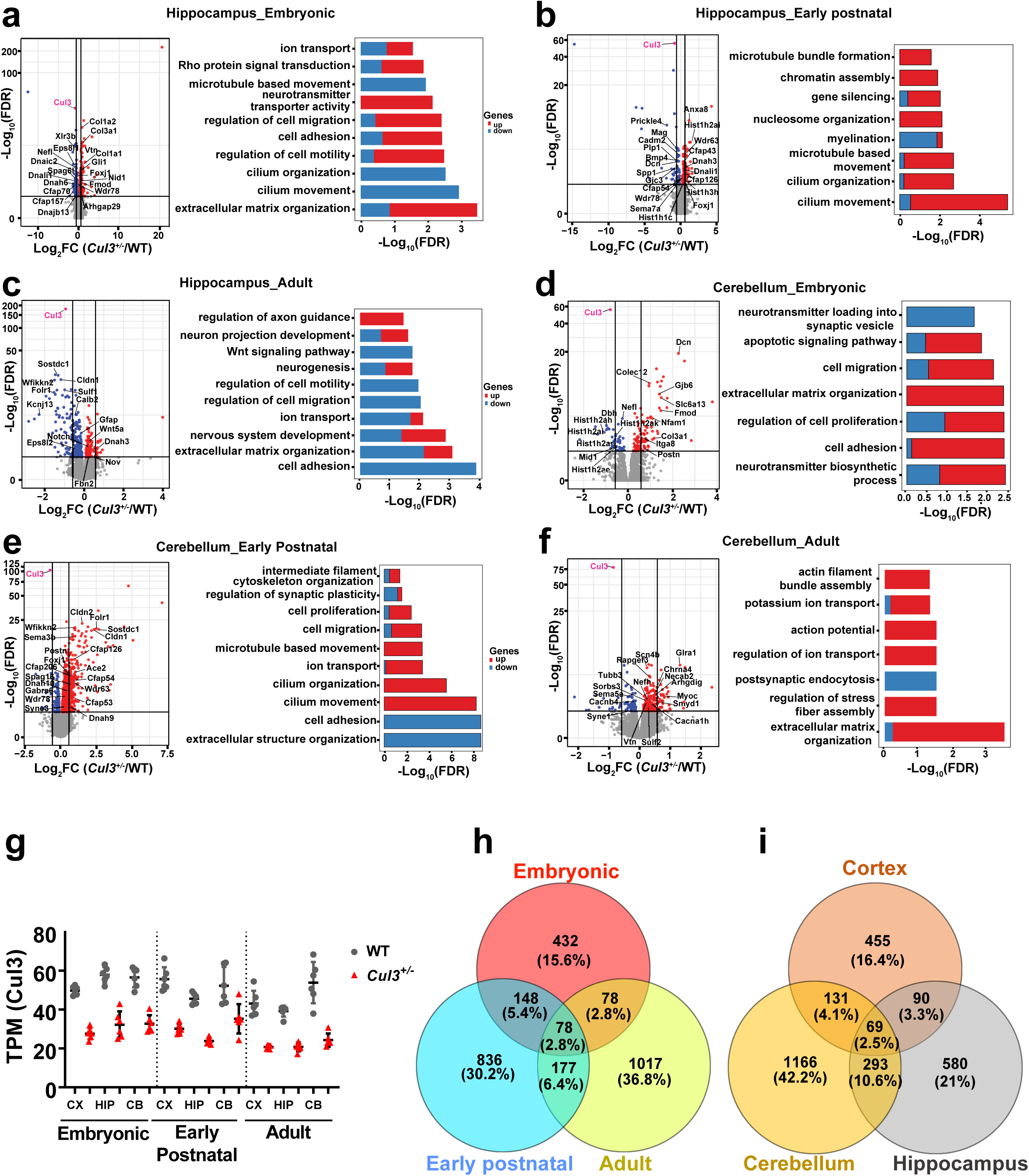
Differential gene expression for individual periods and regions. **(a-f)** (Left) Volcano plots of differentially expressed genes in Cul3^+/-^ *vs* WT for embryonic, early postnatal and adult hippocampus and cerebellum. Genes colored in red are upregulated in *Cul3^+/-^* compared to WT; genes colored in blue are downregulated in *Cul3^+/-^* compared to WT; Cul3 is colored in pink. (Right) GO-terms enrichment of differentially expressed genes by periods and regions. Contribution of up- or down-regulated genes to specific GO terms are shown in blue and red, respectively. (**g**) Dot plot demonstrating TPM value of Cul3 in different brain regions and periods. Dots represent individual animals. (**h**) Venn diagram showing 78 common DEGs across the developmental periods. **(i)** Venn diagram showing 69 common DEGs across brain regions.

**Figure S9.**
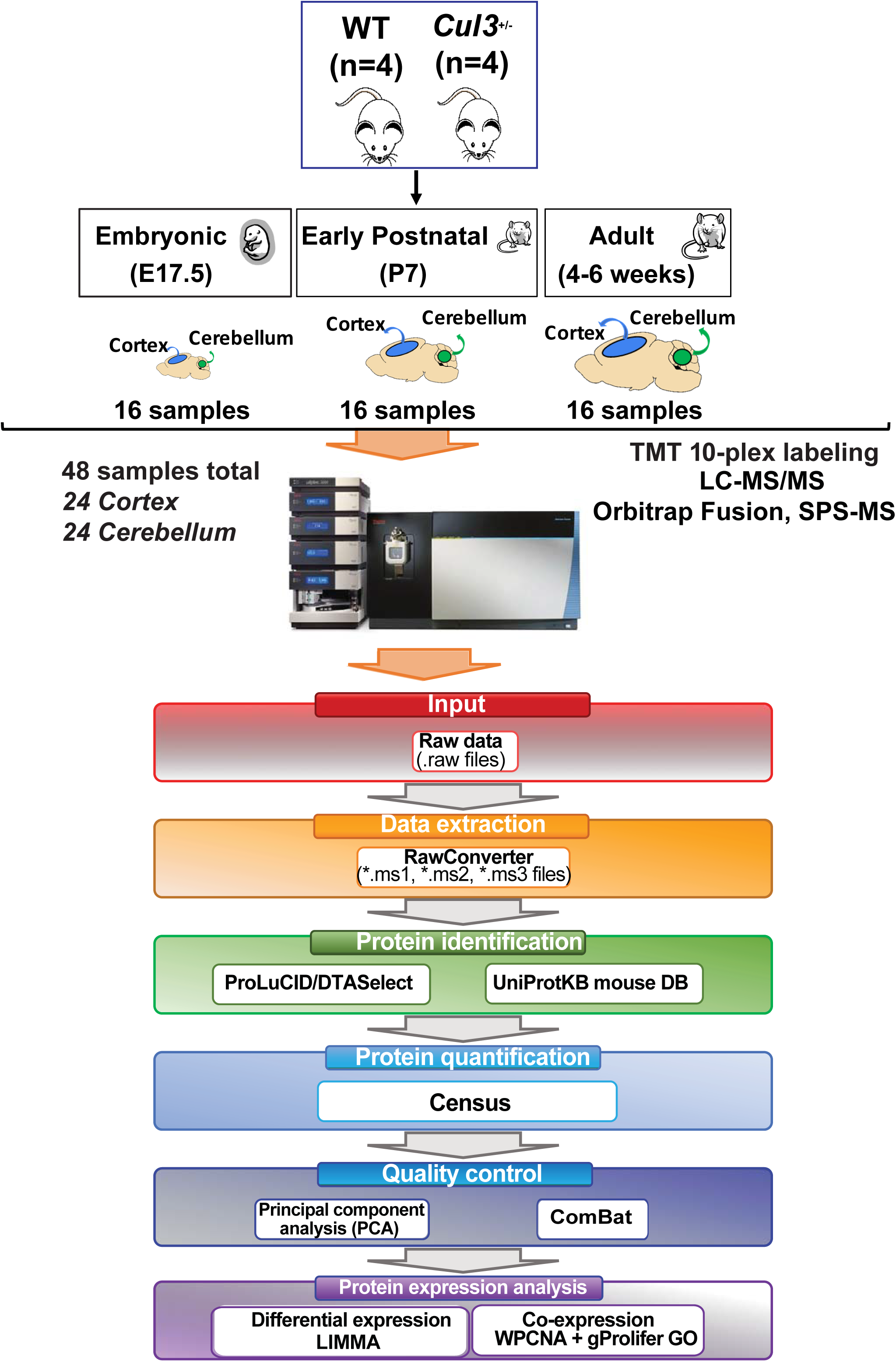
TMT quantitative proteomics experimental design and data analysis workflow. A total of 48 proteomes (24 for each, Cortex and Cerebellum) have been processed in this study by TMT 10-plex labeling followed by LC-MS/MS. Protein Quantification was carried out by Census. Quality control including principal component analyses, ComBat has been performed. LIMMA was implemented for differential protein expression analyses.

**Figure S10.**
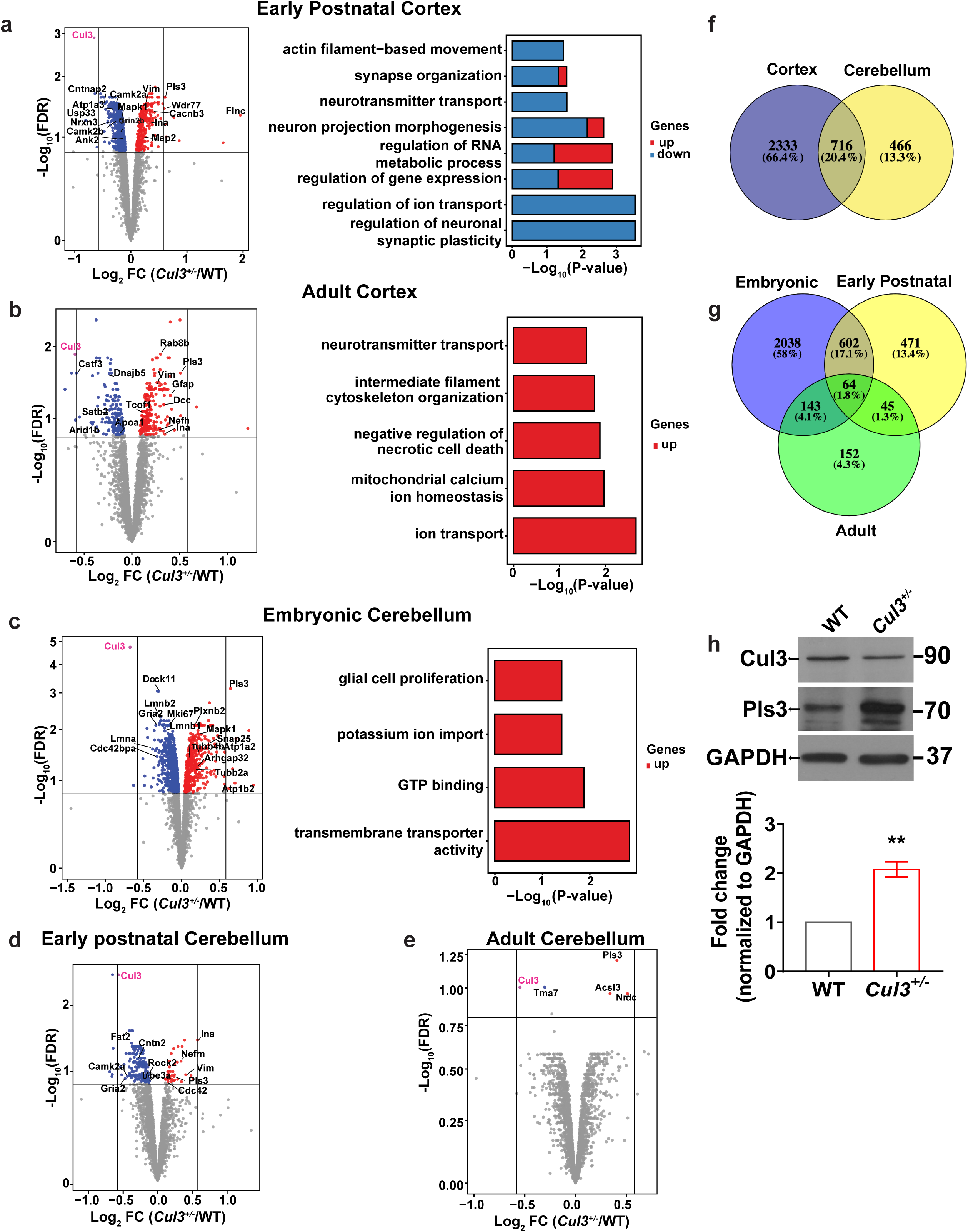
Differential protein expression analyses for individual periods and regions. **(a-e)** Volcano plots of differentially expressed proteins in *Cul3^+/-^ vs* WT (left column) for early postnatal and adult cortex, and embryonic, early postnatal and adult cerebellum. Proteins colored in red are upregulated in *Cul3^+/-^* compared to WT; proteins colored in blue are downregulated in *Cul3^+/-^* compared to WT. (Right) GO-terms enrichment of differentially expressed proteins by regions. Contribution of up- or down-regulated proteins to specific GO terms are shown in blue and red, respectively. Early postnatal and adult cerebellum have no GO-term enrichment. **(f)** Venn diagram showing 716 shared DEPs across brain regions (FDR ≤ 0.15). **(g)** Venn diagram showing 64 shared DEPs across three developmental periods (FDR ≤ 0.15). **(h)** Western blot of Pls3 in embryonic cortex (Upper panel). Densitometry analysis of Western Blot is shown in the bottom panel. Data is represented as mean ± SEM (n=3 per genotype; **p<0.01; two tailed t-Test). Significance above bars represents comparison against WT.

**Figure S11.**
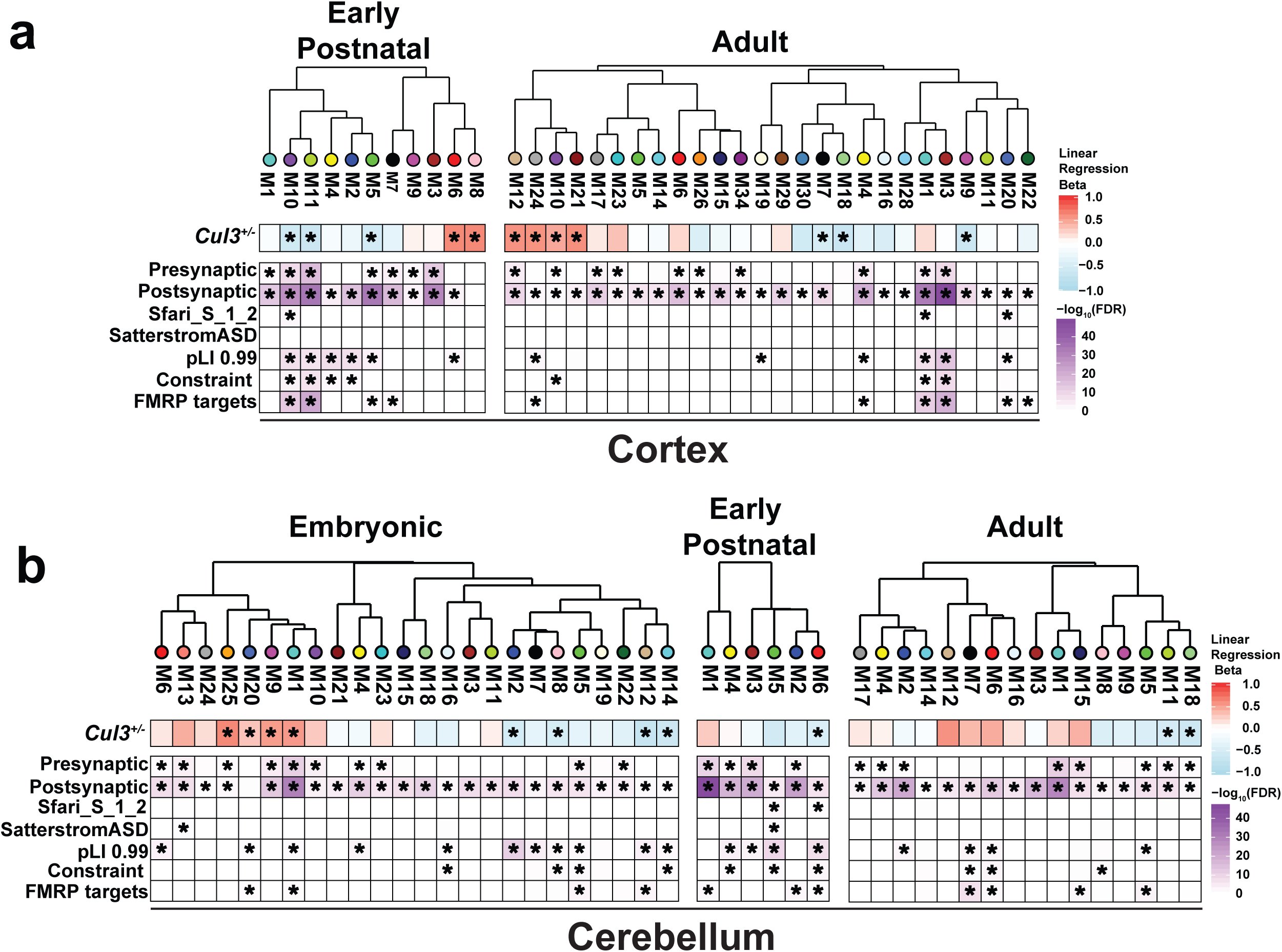
Protein co-expression modules in *Cul3^+/-^* cortex and cerebellum. Hierarchical clustering of protein co-expression modules by module eigengene for *Cul3^+/-^* cortex and cerebellum. Module-genotype associations (*FDR<0.1) are shown below the dendrogram. **(a)** A total of 5 and 7 modules were significantly associated with *Cul3^+/-^* genotype in early postnatal and adult cortex, respectively. **(b)** A total of 8, 1, and 2 modules were significantly associated with *Cul3^+/-^* genotype in embryonic, early postnatal and adult cerebellum, respectively. Module enrichment analyses against literature-curated gene lists with previous evidence for involvement in autism are shown at the bottom (* FDR<0.05).

**Figure S12.**
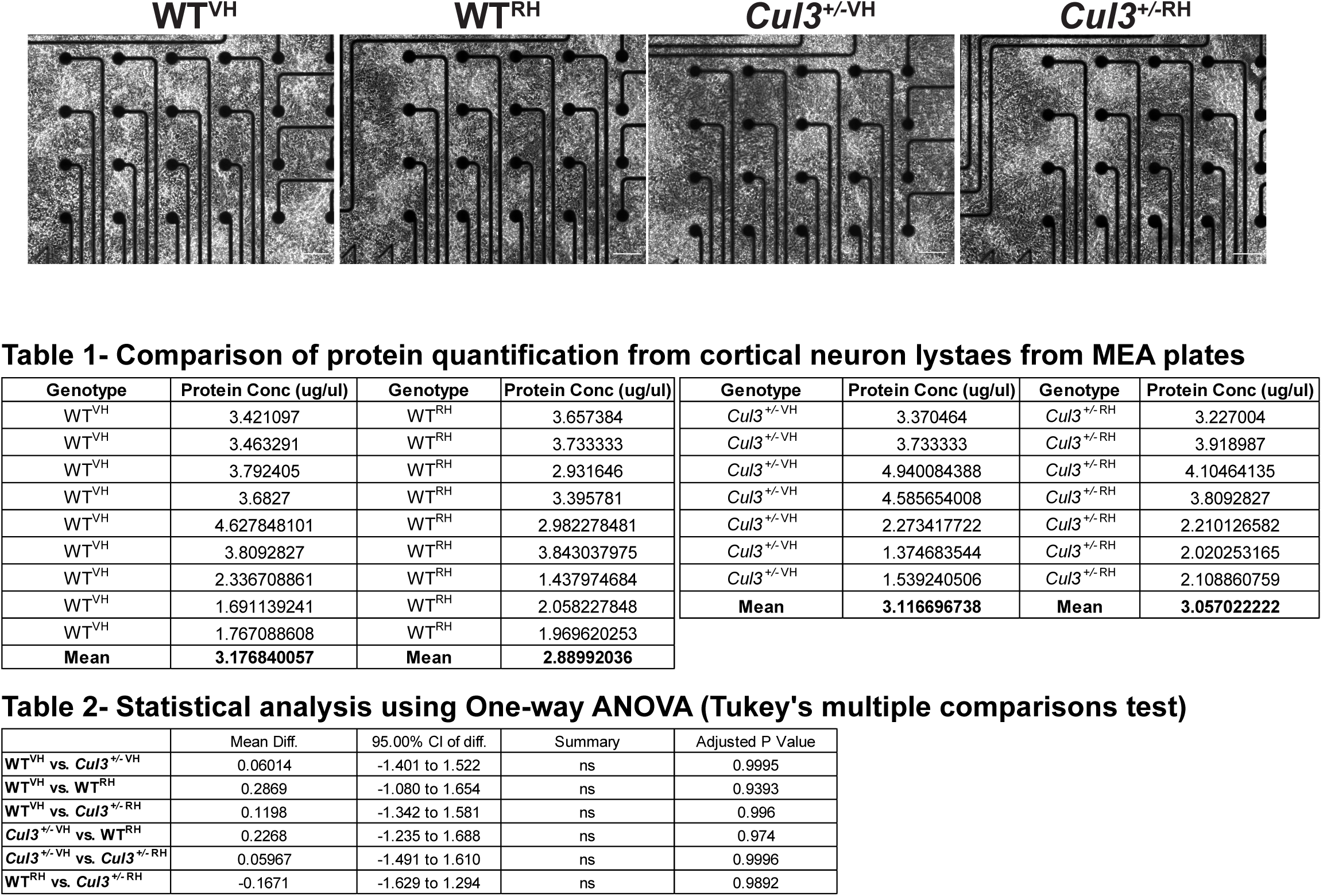
Representative single-well images containing primary cortical neurons surrounding electrodes from the MEA plate for Rhosin-treated WT and *Cul3^+/-^.* Scale bar is 100μm. Table 1 represents protein quantification from WT and *Cul3^+/-^* cortical neurons after completion of MEA recording. Table 2 represents statistical comparison (not significant for each well) of total protein concentration between Rhosin- and Vehicle-treated WT and *Cul3^+/-^* primary neuron culture, suggesting that difference in activity is not due to neuron number.

**Figure S13.**
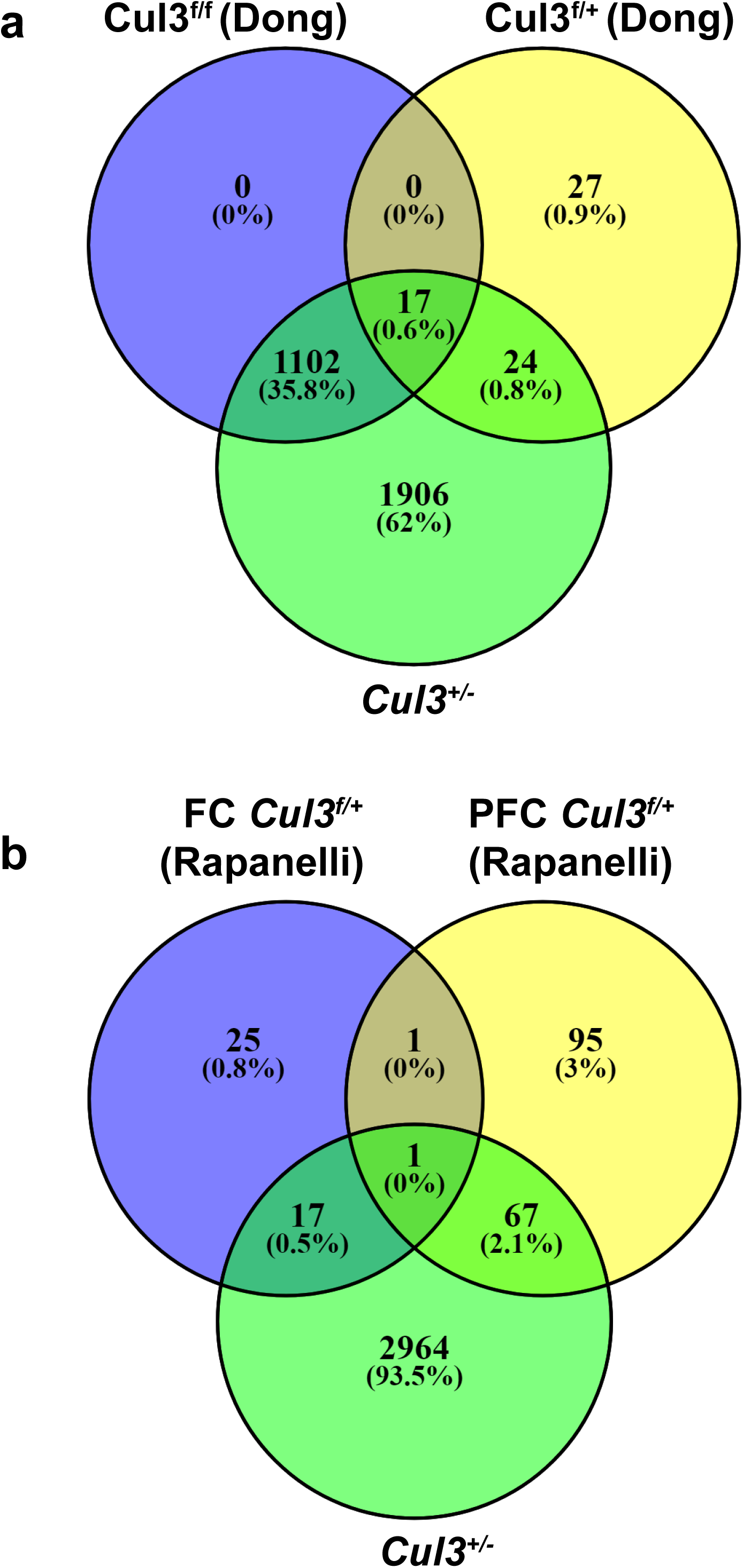
Venn diagram demonstrating the similarity of proteomics results for cortex between two published Cul3 conditional mouse models and our *Cul3^+/-^* CRISPR mouse. The comparison was carried out between nominally significant (without FDR correction) proteins identified in Dong et al. ^31^ and Rapanelli et al. ^29^ with *Cul3^+/-^* CRISPR FDR significant proteins (FDR≤15%). None of proteins identified in Dong or Rapanelli passed FDR significance cut-off, as pointed in the respective studies. **(a)** Comparison of our *Cul3^+/-^* mouse with Dong et al. Cul3^f/f^ Knockout and Cul3^f/+^ heterozygous mice. **(b)** Comparison of our *Cul3^+/-^* mouse with Rapanelli Cul3^f/+^ FC-forebrain specific and Cul3^f/+^ PFC Cul3^f/+^- prefrontal cortex.

**Figure S14.**
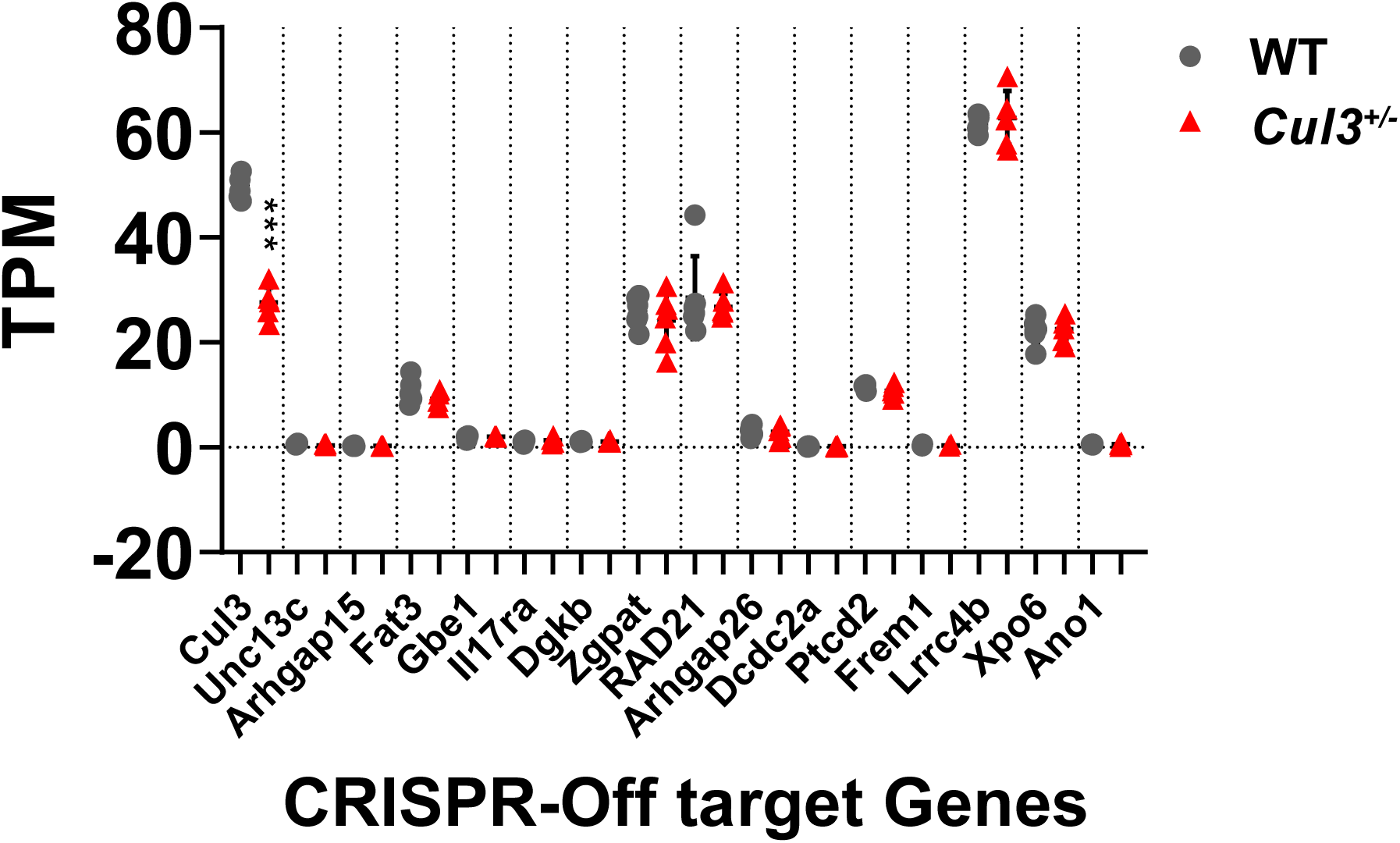
TPM values of CRISPR-off target genes detected by RNAseq. Except Cul3, none of the genes is significantly altered in *Cul3^+/-^* mutant mice. Dots represent individual animals.

## Supplementary Tables titles

**Table S1.** Cul3 mutational spectrum in human carriers with NDDs.

**Table S2.** Brain MRI of *Cul3^+/-^* adult and early postnatal (P7) mouse brain.

**Table S3.** Differentially expressed genes (DEGs) in embryonic, early postnatal and adult cortex, hippocampus and cerebellum.

**Table S4.** Gene Ontology (GO) enrichment analysis of differentially expressed genes.

**Table S5.** Differentially expressed gene (DEGs) clusters used for p-value combination analyses in **Figure 4d**.

**Table S6.** Gene Ontology (GO) enrichment analysis of DEG clusters.

**Table S7.** Differentially expressed proteins (DEPs) in embryonic, early postnatal and adult cortex and cerebellum.

**Table S8.** Gene Ontology (GO) enrichment analysis of differentially expressed proteins.

**Table S9.** Module membership from protein co-expression (WPCNA) analysis in embryonic, early postnatal and adult cortex and cerebellum.

**Table S10.** Gene Ontology (GO) enrichment analysis of protein co-expression modules from WPCNA of cortex.

**Table S11.** Gene Ontology (GO) enrichment analysis of protein co-expression modules from WPCNA of cerebellum.

**Table S12.** CRISPR off-target genes analysis.

**Table S13.** RNA-seq parameters and quality control metrics from Cutadapt, STAR, Picard, and RNA-SeQC for embryonic E17.5, early postnatal P7, and adult brain representing 108 transcriptomes.

